# Continuously fluctuating selection reveals extreme granularity and parallelism of adaptive tracking

**DOI:** 10.1101/2023.10.16.562586

**Authors:** M.C. Bitter, S. Berardi, H. Oken, A. Huynh, P. Schmidt, D.A. Petrov

**Author notes:** Corresponding authors: M.C.B, P.S., and D.A.P.

## Abstract

Temporally fluctuating environmental conditions are a ubiquitous feature of natural habitats. Yet, how finely natural populations adaptively track fluctuating selection pressures via shifts in standing genetic variation is unknown. We generated high-frequency, genome-wide allele frequency data from a genetically diverse population of *Drosophila melanogaster* in extensively replicated field mesocosms from late June to mid-December, a period of ∼12 generations. Adaptation throughout the fundamental ecological phases of population expansion, peak density, and collapse was underpinned by extremely rapid, parallel changes in genomic variation across replicates. Yet, the dominant direction of selection fluctuated repeatedly, even within each of these ecological phases. Comparing patterns of allele frequency change to an independent dataset procured from the same experimental system demonstrated that the targets of selection are predictable across years. In concert, our results reveal fitness-relevance of standing variation that is likely to be masked by inference approaches based on static population sampling, or insufficiently resolved time-series data. We propose such fine-scaled temporally fluctuating selection may be an important force maintaining functional genetic variation in natural populations and an important stochastic force affecting levels of standing genetic variation genome-wide.

## Main Text

Adaptation is a fundamental driver of evolutionary change in natural populations. The field of population genetics has studied this process through the development of models and empirical research that largely assume that most fitness-associated traits are under stabilizing selection and slowly evolve to track an optimal phenotype that changes incrementally through time (1). However, a ubiquitous feature of the habitats in which populations persist is sharp temporal fluctuations in the selective environment. Life has evolved distinct mechanisms to cope with these fluctuations.

At the individual-level, organisms alter their phenotype to track predictable changes in the environment via phenotypic plasticity, or produce a fixed phenotype that minimizes fitness variance amidst unpredictable environmental shifts via bet-hedging (2–5). At the population-level, populations can adapt to different states of the environment via shifts in standing genetic variation, a process termed adaptive tracking (3). What remains unknown is the temporal scale over which populations have the capacity to adaptively track fluctuations in the selective environment. Even in the face of strong, rapidly fluctuating selective pressures, adaptive tracking may be limited if a population lacks genetic variants with sufficiently large selection coefficients to drive differential survival and reproduction at the same pace as the shifting environmental conditions (6–8). Determining whether such conditions are met in natural populations hinges upon well-resolved, time-series genomic data. Procuring such data may ultimately reveal a functional relevance of standing genetic variation that remains hidden from inference approaches using static snapshots of population genomic variation (9–13). Furthermore, such approaches would help clarify the extent to which temporally fluctuating selection could act to maintain adaptive variation in natural populations, an issue of longstanding theoretical and empirical debate (14–19).

Seasonal environments, which fluctuate cyclically and on a timescale that lends to repeated empirical observation, are well-suited to quantify how finely populations adaptively track fluctuating selection pressures. For example, populations of *Drosophila* spp. inhabiting temperate environments expand exponentially from spring through summer as resources become abundant, then crash during the onset of winter as abiotic conditions deteriorate and resources become scarce (20). These boom-and-bust dynamics are a dominant feature of the demographic trends in a range of species, occur on scales shorter and longer than the seasonal scales (21–23), and have been shown to drive the evolution of life-history, reproductive, and stress tolerance traits (24–26).

Investigation into the population genetics of adaptation in seasonally fluctuating environments has a long history (27–30). Recent studies that have leveraged genome-wide sequencing of *D. melanogaster* populations have shown how adaptive tracking keeps pace with fluctuating selection on seasonal timescales (31–33). Notably, Rudman et al. (2022) established that parallel, genome-wide signatures of fluctuating selection can be observed on monthly intervals (∼1-3 generations) across ten replicate populations, suggesting that changes in the direction of selection may occur due to more fine-scale shifts in selective pressures than the most conspicuous environmental fluctuations across seasons (33). Still, the underlying granularity of this process remains unknown: do alleles responding to fluctuating selection over monthly intervals exhibit parallel, deterministic changes on even finer scales in response to the exceptionally dynamic environments in the system, which shift daily across multiple biotic and abiotic axes? Resolving the ultimate limit of adaptive tracking in this system would in turn lend broader insight into the power of temporally fluctuating selection as a process that could preclude the fixation of adaptive alleles, and thus help to maintain variation in natural populations. Furthermore, repeated observation in this system may be leveraged to probe the predictability of the targets of selection across independent bouts of adaptive tracking, an outstanding issue of widespread practical importance (34–36).

Here, we aimed to resolve the scale and pervasiveness of fluctuating selection and the predictability of adaptive tracking via high-frequency monitoring of a genetically diverse population of *D. melanogaster* in an extensively replicated field mesocosm system (Fig. 1). Specifically, we seeded twelve replicate outdoor cages with a genetically diverse population (hereafter, baseline population), generated via four generations outbreeding a panel of inbred reference lines that were originally collected in local Pennsylvania orchards. The experiment spanned from early summer through fall, a period of 10-12 generations (25), during which time the replicate populations were subject to the same general shifts in abiotic and biotic selective pressures across seasons. A generic demographic trend in the system was observed, whereby population sizes expanded rapidly after cage founding, stabilized at a peak density by late summer, and declined during fall (20, 33). Notably, the population trajectories exhibited distinguishable behavior, likely representing unique environmental exposures that alter the particular selective environment within each replicate at any given point in time (Fig. 1). We sampled and quantified patterns of genomic variation weekly between 13 July and 7 September 2021, after which samples were collected on 21 September, 20 October, and 20 December (12 total time points; Fig. 1).

**Figure 1.**
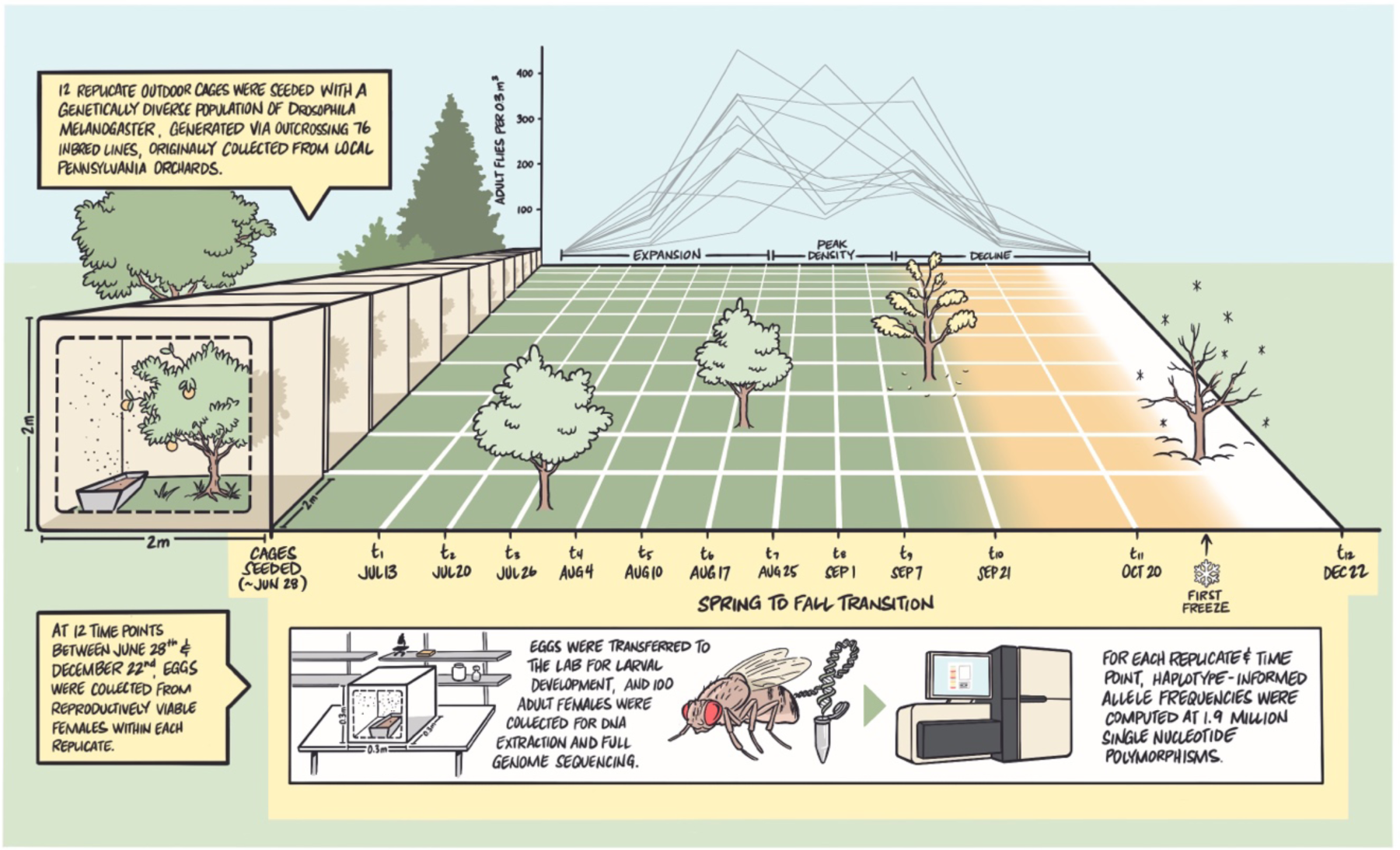
Experimental Schematic. | Twelve replicate outdoor cages in Pennsylvania were seeded with an outbred *D. melanogaster* population, generated via outcrossing 76 inbred lines, originally collected in local orchards. The replicate populations expanded exponentially through mid-summer before declining in early fall, and ultimately crashing following the first freeze in November. At twelve time points throughout the experiment, eggs from the reproductively viable females within each replicate were collected, *in situ*, overnight. The eggs were then transferred to controlled, lab-based conditions for development. One hundred randomly selected female flies from each sample were collected and used for pooled, whole genome sequencing. The resulting sequencing data from each replicate and time point was then used for haplotype-informed allele frequency estimation at 1.9 M single nucleotide polymorphisms previously identified in the inbred reference panel.

The sampling scheme allowed powerful inference of adaptive tracking within the system. Specifically, continuous birth, aging, and mortality of flies throughout the experiment induced a shifting age structure within the replicate populations. As individual viability and fecundity vary as a function of age, it was critical to ensure that our quantification of allele frequencies reflected shifts to the instantaneously reproductive genotypes, or ‘leading edge’, of the population (37). Accordingly, at each time point eggs were collected overnight directly from the mesocosms, and larval development was carried out in lab-based conditions. One hundred adult female flies from the resulting cohort were collected and sequenced in pools, from which we generated high-accuracy, haplotype-informed allele frequencies at 1.9 million single nucleotide polymorphisms (SNPs) (38; see Methods). The resulting data from twelve cages and time points were used to quantify parallel, deterministic shifts in allele frequencies across replicates and infer instances of fluctuations in the selective environment over a range of temporal scales. We additionally compared observed patterns of allele frequency change to an independent study using the same mesocosm system (33) to explore the predictability of independent bouts of adaptive tracking.

### Parallel allele frequency change across summer to fall transition

We aimed to detect whether amidst the inherent complexity of our outdoor study system (Fig. 1), there existed evidence of parallel shifts in genomic variation in response to the shared selective pressures experienced across replicates. We first computed genome-wide F_ST_ between the replicate populations and the baseline to ensure that the evolution of allele frequencies quantified throughout the experiment exceeded sources of biological and technical noise (Fig. S1). Next, we used principal component analysis (PCA) to assess the impact of two key experimental factors, replicate cage and collection time, on variance in allele frequencies throughout the experiment. Segregation of samples along the first principal component (5.54 % variance explained) indicated that variation in allele frequencies across all samples was predominantly driven by collection time point (Fig. 2A-B), a pattern maintained when the analysis is run separately across each chromosomal arm (Fig. S2). The segregation of two replicate cages along the second principal component (Fig. 2A) indicates that background allele frequencies across replicates were potentially perturbed during cage founding (e.g. via bottleneck/drifts or initial idiosyncratic selective events). Yet, these two cages maintain the temporal trend along PC1, further indicating parallel shifts in allele frequencies throughout the experiment (Fig. 2B). An unsupervised clustering (K-means) of the PCA results further validated, over a range of cluster sizes, that the predominant axes of variation in the data were most strongly associated with sample collection time (Fig. S3). In concert, these results suggest that the shared selective pressures experienced by the replicates induced concordant changes in genomic variation throughout the experiment.

**Figure 2.**
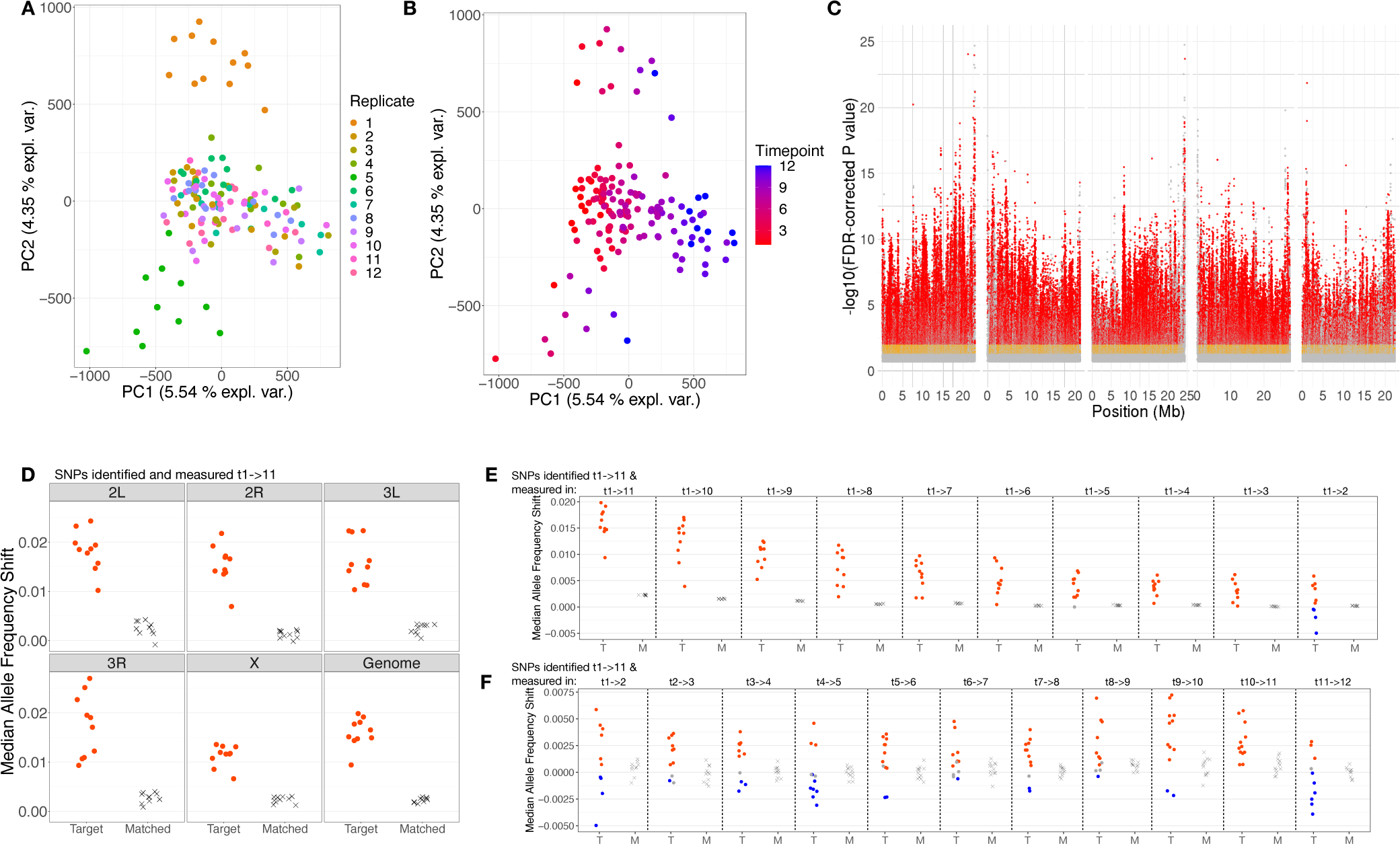
Genome-wide patterns of parallel allele frequency change throughout spring to fall transition. | (A/B) Principal component analysis of normalized allele frequency data (1.9 M SNPs) across all replicates and time points. Sample scores are projected onto the first two principal components and colored by replicate cage number (A) and collection time point (B). (C) Manhattan plot depicting SNPs identified as significant using a generalized linear model and regression of allele frequencies across the entire summer to fall transition (t1->11). Significant SNPs (points) are visualized as a function of genomic position and FDR-corrected P-value, and colored as follows: FDR < 0.2 and effect size > 0.5% (grey); FDR < 0.05 and effect size > 2% (orange); FDR < 0.01 and effect size > 2% (red). (D) Leave-one-out cross-validation of SNPs depicted in (C) whereby the phased frequency dynamics of target SNPs for each left-out replicate are compared to a set of matched control SNPs. Portrayed is the median shift of target and matched control SNPs for each cage, separately for each chromosomal arm and genome-wide. Target site shifts are colored red if the distribution of phased allele frequency shifts was significantly greater, and blue if significantly less, than that of matched control sites (paired t-test, FDR < 0.05). (E/F) Median allele frequency shifts at SNPs depicted in (D), but measured in sequentially shorter intervals (E) and across all pairs of single time points (F). Coloring scheme same to that used in (D).

We next sought to identify the number and genomic distribution of SNPs underpinning parallel changes in genomic variation. Accordingly, we fit allele frequencies to a generalized linear model (GLM; with quasibinomial error variance) to identify SNPs moving systematically across replicates throughout the entire spring to fall transition (time points 1->11; before the total crash of the adult population occurring between time points 11 and 12). This yielded pervasive, genome-wide signal, whereby 28.5% of all SNPs exhibited a Benjamini-Hochberg FDR-corrected P-value < 0.05 (Fig. 2C). Using an algorithm to identify clusters of unlinked loci enriched for significant SNPs, we identified 264 loci underpinning this signal, located across all chromosomal arms and inside and outside known cosmopolitan inversions (33; see Methods) (Supplementary Data File 1). This suggests an oligogenic architecture of adaptive tracking in the system, driven by alleles of relatively large effect size. Specifically, the range of allele frequency shifts for the top SNP of each unlinked cluster (defined as lowest P-value SNP) was 2-11% (median 5.5%), values on par with previous documentation of seasonal allele frequency change in the species (31, 32).

We quantified the degree of parallel, neutral, and/or anti-parallel evolutionary change for individual replicate cages by re-implementing our GLM analysis using a leave-one-out cross-validation approach: sets of GLM significant SNPs were identified iteratively in *n* –1 ‘training’ cages, and the frequency dynamics of the rising alleles (FDR < 0.2, allele frequency change > 0.5 %) were then quantified in the left-out, ‘test’, cage. Each target SNP was matched to a control SNP on the same chromosomal arm and with comparable baseline frequency, recombination rate, and inversion status (see Methods). The distributions of target and control shifts for each cage were then compared to: 1) determine whether the magnitude of target shifts exceeded background allele frequency shifts and therefore the target SNPs exhibited non-neutral dynamics, and 2) infer whether the direction of non-neutral shifts was in a predominantly concordant/parallel direction as that observed in the training cages. This revealed a striking degree of parallelism across all replicates, whereby the median shift of target SNPs within left-out cages was significantly greater than allele frequency change at matched control sites (Fig. 2D). These patterns suggest that even amidst the variation in the genomic composition driven by early founder effects (Fig. 2A), genome-wide allele frequency change in the system exhibited detectably parallel behavior throughout the summer to fall transition.

### Fluctuating selection revealed by higher resolution measurement of allele frequencies

Are alleles exhibiting parallel and systematic changes throughout the summer to fall transition subject to sustained, directional selection throughout this period? Or are fluctuations in the selective environment sufficiently strong to drive reversals in the direction of frequency change on finer timescales? To explore if the direction of selection on t1->11 SNPs was constant throughout the experiment, we first quantified allele frequency shifts of each set of t1-

>11 target SNPs (identified above; Fig. 2C/D) in sequentially shorter intervals. In this way we explored whether there exists a limit at which replicates no longer exhibited deterministic shifts in allele frequency (the magnitude of phased allele frequency shifts no longer differed from matched control shifts) and/or whether evidence of anti-parallel behavior emerged on shorter intervals (significantly negative, median phased shifts at target SNPs), indicating fluctuating selection for the variants that systematically moved in a consistent direction from the beginning to the end of the experiment. As the dynamics of each set of target SNPs were measured using the left-out cage, inferred changes in the direction of SNP movement are insensitive to statistical artifacts, such as regression to the mean.

We observed persistently deterministic, and parallel behavior of target SNPs from the longest interval (t1-> 11; 15 weeks), to the two-week interval (t1->3) (Fig. 2E). The magnitude of the allele frequency change declined monotonically with test interval length. Strikingly, our shortest interval length (1 week, t1->2) continues to indicate predominantly deterministic shifts at target SNPs, but also provides us with the first evidence of anti-parallel behavior across replicates (both red and blue points observed within measurement interval; Fig. 2E). Figure 2F expands the set of test intervals to include all pairs of single time points (t1->2, t2->3…t11->12), the first nine of which are weekly intervals. Concordant with the t1->2 comparison, we see largely deterministic shifts at target sites in left-out cages, yet both parallel and anti-parallel behavior. This indicates that even those SNPs inferred to shift systematically and in parallel throughout the summer to fall transition (Fig. 2D), were in fact subject to numerous bouts of fluctuating selection on much finer temporal scales.

### Continuously fluctuating selection underpins rapid adaptive tracking

How many times does the direction of selection fluctuate throughout the experiment? Most cages experienced at least one reversal in the direction of selection on SNPs identified from t1->11 (Fig. 2F). We hypothesized that sets of alleles identified on shorter intervals, with negligible net change throughout the experiment, may reveal even greater incidence of fluctuating selection in the system. We first defined the shortest interval length over which we had power to detect SNPs shifting systematically, and in parallel, across replicate cages using GLM (Fig. S4). The models regressing allele frequencies across five time point intervals (e.g. t1->6, t2->7…t7->12) represented the shortest interval length in which all comparisons yielded pervasive, genome-wide significant and large effect size variants across all chromosomal arms (Fig S5-13). We thus focused on a temporal interval length of five time points to explore the complex selective environment observed in Figure 2F, but in the supplement also include results from all interval lengths and comparisons to ensure that observed trends were not simply a feature of SNPs identified on this timescale. Furthermore, for this analysis we discarded those SNPs identified as strongly parallel across the t1->11 interval (GLM FDR < 0.05; ∼28% of all SNPs), as these exhibit predominantly directional change (Fig. 2E) and may mask the finer-scale fluctuations in the selective environment we ultimately aimed to uncover.

We first used leave-one-out to quantify the behavior of SNPs discovered for all five time point intervals across the time points over which they were identified. In accordance with SNPs identified from t1->11, those identified on five time point intervals exhibited largely parallel shifts across replicates: for the majority of left-out cages and intervals, the magnitude of frequency shifts at target SNPs were significantly greater than those at matched control SNPs (red points, Fig. 3A). Notably, during the first two intervals (t1->6 and t2->7) two cages (replicates 1 and 5) exhibited predominantly anti-parallel allele frequency change (blue dots). These are the same cages offset along PC2 in Figure 2A/B, and this dynamic likely provides further illustration of perturbations in genome-wide allele frequencies (via either selection or bottlenecking/drift) can drive idiosyncratic behavior across replicates in the early season that are ultimately overcome by parallel shifts in genomic variation once population sizes become sufficiently large (Fig. 1).

**Figure 3.**
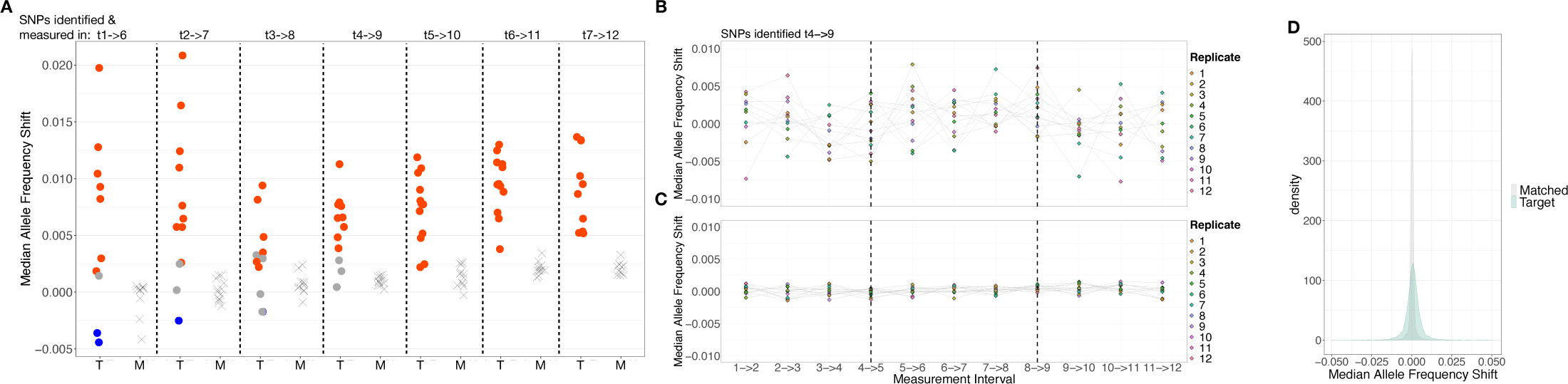
Pervasiveness of fluctuating selection. | (A) Median allele frequency shift for target (T) and matched control SNPs (M) for each left-out cage, using sets of SNPs identified and measured within each five time point interval. Median shifts for target SNPs are colored red if the distribution of phased allele frequency shifts was significantly greater, and blue if significantly less, than that for matched control sites (paired t-test, FDR < 0.05). (B/C) Median allele frequency shifts in each left-out cage (as in A) measured within each pair of single time points for sets target SNPs (B) and matched control SNPs (C) identified t4->9. (D) The distribution of median allele frequency shifts for sets of target (green bars) and matched control SNPs (grey bars) derived from all five time point intervals and measured across pairs of single time points (variance of distributions is significantly different; F-test, F = 0.17, P value < 0.001).

Next, we quantified the number of times fluctuating selection produced changes in the inferred direction of selection acting on replicate populations throughout the experimental period. Specifically, we measured the shifts of target SNPs and matched controls identified on five time point intervals across all pairs of single time points (i.e., t1->2, t2->3…t11->12; as in Figure 2F). Figure 3B-C depicts the results of this analysis for sets of SNPs identified from t4->9: points are colored corresponding to replicate cage and represent the median shift of test SNPs (B) or matched control SNPs (C), with grey lines connecting the shifts for each cage across measurement windows. Across replicates, the median number of flips in the dominant direction of selection was five (flips inferred when shifts at test SNPs changed from being significantly greater to significantly less than matched controls, or vice versa). This indicates that selection coefficients of selected alleles repeatedly flip in sign from positive to negative over weekly timescales, and that the replicate populations adaptively track fluctuations in the environment that occur over much finer timescales than the dominant ecological phases (e.g., population expansion, peak density, or decline; Fig. 1). The underlying magnitude of median allele frequency change at target SNPs throughout these flips was frequently greater than 1% (Fig. 3D), corresponding to selection coefficients that can exceed 10 % per generation, and thus indicating exceptionally strong fluctuating selection on this timescale. Our leave-one-out approach and comparison to matched controls ensures these fluctuations are not a product of regression to the mean, or that the identified SNPs simply exhibit stochastic/noisy behavior due to their intrinsic characteristics (e.g., starting frequency, recombination rate, or inversion status). Qualitatively similar dynamics were observed across all interval lengths and possible comparisons (Fig. S16-23).

### Predictable patterns of genomic variation underpin adaptive tracking

Our results thus far indicate that even amidst differences in starting allele frequencies (Fig. 2A/B), adaptive tracking across the twelve replicate cages proceeded via detectably parallel shifts in genome-wide variation (Fig. 2D). Similarly, using the same mesocosm system, a largely overlapping set of founder lines, and monthly allele frequency estimation between July and November of 2014, Rudman et al. (2022) identified parallel allele frequency change across ten replicate populations. This independent dataset from the same system provides the unique opportunity to ask: how predictable are the genomic underpinnings of adaptive tracking across independent bouts of evolution in cyclically fluctuating environments? Across both experimental years, the replicate cages exhibited boom-and-bust dynamics and were exposed to the same generic seasonal shifts in abiotic conditions and showed parallel genomic shifts across the cages (33). However, it is possible that differences in starting allele frequencies, in combination with year-specific environmental conditions, drive largely unpredictable patterns of genomic variation from year to year. While there is substantial evidence for predictable phenotypic trajectories in response to repeated selective regimes in this and other systems (25, 39), evidence of predictability at the genetic level is largely confined to traits with simple genetic bases (40–42) (but see (43)).

We tested whether independent bouts of adaptive tracking were predictable by comparing patterns of allele frequency change quantified here to those reported by Rudman et al (2022). We first explored genome-wide patterns of differentiation across all samples collected by each study using PCA. While the clustering of samples by year along the first principal component indicated strong differentiation of genome-wide allele frequencies between studies, the separation of early from late season samples along the second principal component suggested that changes in allele frequencies through time may have been similar across years (Figure 4A). To test this systematically, we computed Pearson’s correlations between genome wide allele frequency shifts from all possible intervals of the current study (all two to eleven time point intervals) and each monthly interval of the Rudman et al. study. These correlations provided a directional metric of concordance, whereby positive correlations indicated that genome-wide allele frequency changes were predominantly in the same direction between the tested 2014 and 2021 intervals, and negative correlations to genome-wide SNPs exhibiting predominantly anti-parallel behavior across compared intervals. To infer whether the magnitude of observed correlations exceeded those expected by chance, we computed a null distribution of correlations by randomly shuffling the frequency changes for each comparison across SNPs (values shuffled between SNPs matched by chromosomal arm, starting frequency, recombination rate, and inversion status) and recomputed each comparison’s correlation. The distributions of empirical correlations exhibited an enrichment of comparisons that were both significantly correlated and anti-correlated between years (Fig. 4B; F-test, F = 3821, P << 0.001), a pattern that is maintained when running this analysis separately across each chromosomal arm (Fig. S24). This enrichment of elevated correlations indicates that the same loci are repeatedly subject to selection across years.

**Figure 4.**
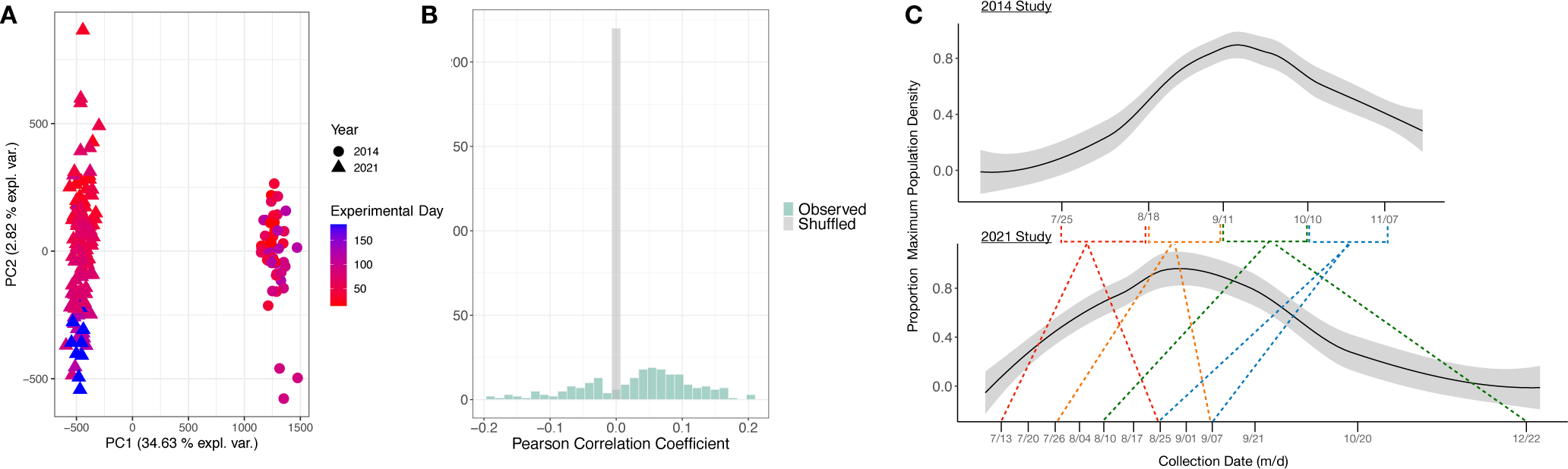
Predictable patterns of selection across independent bouts of adaptive tracking. | (A) PCA of all samples collected during monitoring of adaptive tracking within the mesocosms in 2014 and 2021, colored by experimental day (days since replicate cage founding). (B) Histogram distribution of Pearson correlations computed between genome-wide allele frequency shifts for each monthly interval of the 2014 experiment (N = 4), and each interval of the 2021 experiment (N = 56). Bar color correspond to correlation coefficients generated via site information permutations (grey) and observed values (green). (C) Mesocosm-wide population demographic dynamics for the 2014 and 2021 experiments, whereby mean population size is standardized to maximum observed population density. Dashed lines connect each monthly interval of the 2014 experiment to the best matched interval of the current study, based on genome-wide Pearson correlation between allele frequency shifts.

To explore whether those intervals with heightened allele frequency concordance across years aligned with conspicuous shifts in the selective environment, we identified the interval of the current study that was best matched (i.e., highest genome-wide correlation of allele frequency shifts) to each of the four monthly intervals of the 2014 experiment. Strikingly, the first three monthly intervals of the 2014 experiment were best matched to the t1->7, t3->9, t5->12 intervals of this study, generally overlapping with the periods of population expansion, peak density, and decline (Fig. 4C). The final monthly interval was best matched to T7->9, another mid-late season period, but we note that the correlation coefficient for this matching was an order of magnitude less than those for the first three monthly intervals (though still significant based on our empirically-derived null distribution). Ultimately, these matchings suggest that amidst stochasticity in year-to-year abiotic conditions, the cyclical boom and bust dynamics of the populations induce a repeatable ecological filter that shapes patterns of selection that recurrently affects the same set of loci. Such patterns generally conform to long-standing ecological hypotheses regarding the distinct selective environments associated with population expansion and contraction and corroborate mounting evidence of pervasive, genome-wide selection throughout the progression of seasonal adaptation (21, 27–29, 31–33).

## Discussion

The pervasive fluctuating selection we report here reveals key insights into the dynamics of adaptive tracking, with implications for patterns of genome-wide diversity and the maintenance of genetic variation in natural populations. Specifically, we identified changes in the dominant direction of selection repeatedly over multiple weekly intervals throughout the course of the experiment (Fig. 3). This indicates that our outbred population harbors alleles at relatively high frequency that exhibit large selection coefficients that flip sign in accordance with fluctuations in environmental conditions over exceptionally short timescales. More specifically, across the time points over which each instance of fluctuating selection was inferred, the environment must have changed in such a way to induce trade-offs in the combination of traits underpinned by these alleles. Thus, even within each of the dominant ecological phases (e.g., population expansion), the adaptive direction of multi-variate phenotypic change likely shifts constantly as changing population densities interact with specific combinations of abiotic conditions. Furthermore, it is likely that those alleles with repeated sign flips exhibit antagonistically pleiotropic effects, whereby they confer advantageous values for the specific traits most readily impacting fitness during intervals in which they are favored, but induce disadvantageous trait values for the sets of traits subject to selection during intervals in which they are selected against (44–46). As these alleles exhibited undetectable systematic change throughout the experiment, they would likely be inferred to be neutral using inference approaches based on coarse temporal sampling or static snapshots of population genomic data. Thus, our high frequency sampling revealed a functional relevance to standing variation that would have remained hidden, bolstering recent calls for the procurement of well-resolved, time series genomic sampling (9, 10, 12, 13, 31–33).

The patterns of fluctuating selection we observed are likely to have appreciable impacts on genome-wide patterns of diversity within the species. Specifically, selection in various forms (e.g. background selection, selective sweeps, fluctuating selection) have been shown to broadly reduce diversity at linked neutral sites (47–58). In the case of temporally varying selection, it has been shown that even modest fluctuations can have substantial impacts on genome-wide patterns of diversity (58). Thus, the strong, rapid, and genome-wide signals of fluctuating selection we quantified are likely to alter patterns of diversity in a pronounced manner. This holds implications for recent debate as to whether elevated selection in large population-size species can help resolve ‘Lewontin’s Paradox’ – the puzzling observation that neutral genetic variation does not scale with census population size as expected (58–62). While some studies have inferred that selection alone is insufficient to explain the narrow range of polymorphism observed across species (59, 60), the exceptionally strong selection quantified here calls this assertion into question and warrants increased effort towards the procurement of well-resolved time-series genomic data in other systems.

Our results may additionally lend insight to the mechanisms maintaining variation in natural populations. For example, a large body of theoretical models has suggested that temporally fluctuating selection pressures are not likely to maintain balanced polymorphisms for genotypes specialized to different states of the environment, but rather drive the fixation of optimally plastic genotypes that are able to alter their phenotypes to match the shifting conditions (3, 17, 63, 64). Such research has led to the viewpoint that spatially varying selection is the dominant driver of balanced polymorphisms, and thus the maintenance of variation, across species’ ranges (18, 65). Yet, more recent models have begun to incorporate various forms of multi-locus dominance, particularly environmentally-induced reversals of dominance, and demonstrated that polymorphism at a large number of loci can be maintained in temporally fluctuating environments without imposing unsustainable genetic load on the population (15, 19, 44, 66). Our observations add credence to these models, demonstrating how rapidly fluctuating selection could limit the fixation of adaptive alleles in a population. But whether it indeed is the key force maintaining variation across species ranges is an outstanding empirical issue.

It is critical to note that our detection of rapidly fluctuating selection was likely contingent upon our strategy for quantifying changes in genomic variation in the system. Specifically, changes in allele frequencies over our shortest sampling intervals likely represent subtle, yet significant, shifts in the relative contribution of specific genotypes to the next cohort of flies. Our detection of such fine-scale shifts hinged upon our highly replicated system and standardized sampling of the instantaneously reproductive population at each collection point. Furthermore, as demonstrated in Figure 2D/E, the resolution of sampling throughout the season dictates our ability to detect instances of fluctuating selection, which are averaged out and remain hidden when evolutionary change is quantified over longer timescales (13, 67, 68). We thus conclude that the relatively sparse empirical evidence of fluctuating selection across systems may be due to an ascertainment bias of quantifying evolutionary change on longer timescales (33, 69). Indeed, research into patterns of spatially varying selection has indicated that signals of balancing selection over exceptionally fine spatial scales (‘microgeographic’ adaptation) can be masked by coarse sampling (70). Thus, this pattern may be recapitulated under instances of temporally varying selection, though the ubiquity of this dynamic across study systems and species remains to be determined.

Our findings additionally add to a growing body of research into whether adaptation across independently evolving replicate populations proceeds via predictable shifts in genomic variation. While numerous observations of predictable phenotypic evolution are reported in this and other systems (25, 39), signals of genomic predictability are largely confined to traits with relatively simple genetic bases (e.g., 40–42). Here, we detect a large degree of within-year parallelism and across-year predictability, despite hundreds of putatively unlinked loci driving patterns of adaptive tracking in the system (Fig. 2C). Strikingly, this parallelism and predictability is evident despite substantial variance in starting allele frequencies across replicate cages within and across years (Fig. 2A/B; Fig. 4). This demonstrates the oligogenic architecture of adaptive tracking in the system is resilient to perturbations of background/starting allele frequencies, likely a result of the additivity of the marginal effects of alleles underpinning the evolving traits and the very strong selective pressures in the system (71).

The predictable patterns of genomic variation across study years further suggests that despite variation in specific environmental conditions, there exists a generality to the suite of selective pressures imposed on the populations. We hypothesize this generality is in large part driven by the boom-and-bust ecological dynamics observed in our mesocosm populations, which is ubiquitous to natural populations of *Drosophila* spp. as well as range of species with differing life-history characteristics (20–23). This hypothesis is bolstered by the fact that the most concordant periods of allele frequency change across years were those when the population sizes were in similar phases of the boom-and-bust cycle (Fig. 4D), as well as previous studies reporting concordant allele frequency shifts between the spring and fall over multiple years (31) and across widely dispersed *D. melanogaster* populations (32). It is not feasible to disentangle the relative contribution of these biotic forces from covarying abiotic selective pressures (e.g., temperature) with the data collected here, though may be possible in the future through a systematic manipulation of these variables, such as a staggered initiation of the replicate cages and population trajectories.

In conclusion, the allele frequency changes we observed illustrate an instance of adaptive tracking underpinned by strong, rapidly fluctuating selection that induces parallel allele frequency change across twelve independent replicates. Such dynamic patterns of selection are likely to have pronounced impacts on patterns of genetic diversity within the species and may act to maintain variation across the species’ range. Furthermore, given that the observed adaptive tracking is likely driven by continual selection and reshuffling of standing variation, it is not implausible that we could detect evolutionary change over weekly, sub-generational, intervals. Indeed, previous studies have quantified selection in the wild occurring on similar timescales, though these are largely confined to pulse selective events, such as hurricanes (72), winter storms (73), harvest and poaching (74), or heat waves (75).

Thus, while this may be the first instance of sustained evolutionary change at this tempo, the phenomenon may in fact be widespread in populations evolving with overlapping generations and revealed by sufficiently resolved time-series genomic data. Finally, the dynamic nature of adaptive tracking observed lends promise towards the capacity for species to adaptively track the changes in the environment induced by global change. Still, an important consideration is how dependent such fine-grained adaptive tracking is on the total amount of genetic variation present in populations, which itself may be reduced as global change progresses (76, 77).

Ultimately, resolving the underlying causal genes underpinning adaptive tracking will lend insights into the total number of loci necessary for fine-grained, rapid adaptation in complex selective regimes.

## Methods

### Study population and mesocosm system

We quantified and tracked shifts in genomic variation of a genetically diverse *Drosophila melanogaster* population over a six-month period in a highly replicated, field mesocosm in Philadelphia, Pennsylvania. The study population was derived via outbreeding a panel of 76 inbred strains originally collected wild from Linvilla Orchards, Media, PA (24). Each line had been previously sequenced to a minimum genome-wide coverage of 50x, following 20 generations of full-sib mating (as described in (38)). An outbred, baseline population was initiated via combining 10 males and 10 females from each line into each of four, 30 cm^3^ indoor cages. Each subpopulation was subject to four generations of unmanipulated recombination and population expansion, a process aimed at purging deleterious alleles that likely accumulated throughout inbreeding. Eggs from the fourth generation were collected and density controlled in vials. Upon eclosure, 500 randomly selected males and females were released into each of twelve two m^3^, outdoor mesh cages.

Each replicate cage harbored a single dwarf peach tree (non-fruiting) and was exposed to natural environmental conditions and insect and microbial communities (33, 78). Four hundred ml of Drosophila media (‘Spradling cornmeal recipe’), provided in 900 cm^3^ aluminum loaf pans, was placed in a shaded shelving unit within each cage three times per week, providing the only source of food and egg laying substrate. Egg laying upon each loaf pan was carried out for two days, after which the pan was enclosed with a mesh lid to prevent any further laying. The pan was then monitored daily, and the lid removed after eclosure was first observed. This process generated a near continual input of new flies into each replicate cage, generating evolution with overlapping generations and a population age-structure that varied throughout the duration of the experiment. The census size of the adult population was estimated five times throughout the progression of the experiment (19 July, 5 August, 20 August, 17 September, and 20 October). To estimate population sizes, we photographed four, 0.3 m^3^ transects on the ceiling of each mesocosm within 30 minutes of sunsets (when adult flies were aggregated on the mesocosm walls). The number of flies within each transect were counted using semi-automated image processing, the average of which was used as the estimated number of adult flies per m^3^ for a given cage and time point.

### Sample collection, sequencing, and allele frequency estimation

We conducted high-frequency sampling of the cage replicates to quantify patterns of genomic variation throughout the experiment. The first sampling was conducted on 13 July 2021, three weeks after the 21 June seeding of the mesocosms (a period aiming to further purge residual deleterious mutations and allow the lab-adapted population to experience 1-2 generations of field selection). Samples were collected weekly for the first 9 weeks of the experiment (13 July – 7 September), after which samples were collected on 21 September, 20 October, and following the first freeze and population crash on 20 December (12 total time points; Fig. 1). Note that egg samples for two replicates (replicate 3 and 7) were lost during the t1 sampling point and eggs for one replicate (replicate 5) were lost during the t3 sampling point. As a consequence, allele frequency data is missing for these samples/time points.

Throughout the experiment, the replicate populations evolved with overlapping generations and thus exhibited shifting age-structure as the experiment progressed (Charlesworth 1980/1994). It was thus critical to account for this demographic structure in quantifying the ecological and genetic state of the system to ensure that our quantification of allele frequencies reflected shifts to the instantaneously reproductive genotypes, or ‘leading edge’, of the population (37). Accordingly, at each time point eggs were collected overnight directly from the mesocosms, and larval development was carried out in lab-based conditions, where eggs developed and eclosed to F1 adults in 30 cm^3^ cages. 3-5 days post-eclosure, a random set of 100 females were sampled from each cage and preserved in 99% ethanol at –20° C for later DNA extraction.

We collected an additional, independent sample of 100 females from each cage at time points 3 (26 July) and 10 (20 October) to quantify biological variation in our allele frequency estimates. Furthermore, an additional four replicate samples of the baseline population prior to field exposure were preserved for genomic analysis. Finally, it is important to note that the final sample point (22 December) occurred following the crash of the adult population, and thus sampling methods for this time point were altered such that larvae were collected directly from the remaining loaf pans within each cage (i.e., no reproductively viable adults remained in the cage at this time point, but larvae persisting in cage loaf pans were taken into the lab and reared to adults for pooled sequencing). Additionally, two replicate mesocosms were unable to be sampled at this final time point, reducing replication. We accommodate for these differences in sampling of t12 by focusing analysis on patterns of variation throughout the summer to fall transition (time points 1 through 11; see below).

We generated high-accuracy, haplotype-informed allele frequency estimates at most of the common polymorphic sites within the species. Specifically, genomic DNA was extracted from each homogenized pool of 100 flies using the Monarch Genomic DNA Purification Kit (New England Biolabs). DNA quality was assessed using Nanodrop and quantified with Qubit, before whole-genome sequencing libraries were generated for each pooled sample using the Illumina DNA Prep Tagmentation Kit. All samples were sequenced on Illumina Novaseq 6000 flow cells using 150 bp, paired-end reads. A total of three separate sequencing rounds were required to generate sufficient coverage across all samples, and technical replication across rounds were used to quantify the negligible impact of batch effects on allele frequency estimation (Fig. S1). Raw sequencing reads were trimmed of adapter sequences and bases with quality score < 20, and aligned to the *Drosophila melanogaster* v5.39 reference genome using bwa and default parameters (79). Aligned reads were deduplicated using Picard tools (http://broadinstitute.github.io/picard/) and the final set of reads for each sample was down-sampled to obtain an equivalent, genome-wide coverage of 8x across all samples.

We computed haplotype-derived allele frequency estimates using a local inference method and pipeline developed by Tilk et al. (2019). Briefly, using an expectation-maximization algorithm for estimating haplotype frequencies in a pooled sample from mapped sequence reads (originally developed by (81)), we obtained allele frequencies at sites previously identified in the genome sequences of each founding inbred strain. We conducted haplotype inference in window sizes that varied proportionally to the length of un-recombined haplotype blocks expected as a function of the estimated number of generations since the original outbreeding. Allele frequencies were compiled to only include those sites with an average minor allele frequency > 0.02 in the baseline population, and present in at least one evolved sample at a MAF > 0.01. See Tilk et al. 2019 for validation that this method produces an accuracy of allele frequencies at a genome-wide depth of 5x comparable to those obtained from raw allele frequencies generated via standard pool-seq data and 100x coverage. *Quantifying genome-wide patterns of variation*

All statistical analysis of allele frequency data was carried out in R v. 3.5.6. We used a population genetic summary statistic, F_ST_, to determine whether observed shifts in genomic variation exceeded sources of technical and biological noise throughout the sampling period. Specifically, we quantified genomic differentiation through time by computing F_ST_ between each evolved sample and each of the four baseline samples (F_ST_ was computed separately for each polymorphic site and averaged across all sites to obtain a genome-wide mean of differentiation). Next, we quantified sources of biological and technical noise by computing F_ST_ pairwise between technical sequencing replicates (same 100 flies and pooled DNA extract, different library preparation and/or sequencing rounds; N = 20 pairwise comparisons) and biological replicates (same cage/time point, different sample of 100 flies; N = 17 pairwise comparisons). We tested whether evolutionary-driven changes in genomic variation exceeded sources of noise using a paired t-test, and whether evolved samples became increasingly differentiated from the baseline population throughout the experiment using a generalized linear model (family = gaussian; link = logistic).

We tested whether variance in allele frequencies across all collected samples was dominated by replicate cage or collection time point using principal component analysis (PCA) (*prcomp* package). Allele frequencies were centered and scaled, and each sample was projected upon the first two principal components. We ran this analysis both genome-wide, as well as separately for SNPs on each chromosomal arm. We conducted K-means clustering of each sample’s PC scores (*stats* package) to, in an unsupervised manner, quantify the extent to which PC clustering was predominantly driven by replicate cage or collection time. Specifically, if K-means clustering of sample points corresponded to sample collection time point (i.e., segregated early from late season samples), this provided a quantitative validation of trends visualized along PC1 and PC2.

### Target SNP identification and characterization of genetic architecture

We identified single nucleotide polymorphism (SNPs) shifting systematically through time and across replicates by fitting allele frequencies to a generalized linear model (formula: allele frequency ∼ time point) (33, 81). Allele frequencies were weighted by the total number of chromosomes sequenced (N = 200) and depth per sample (38). GLMs for each site were run with a quasibinomial error model, and P-values were adjusted using the Benjamini-Hochberg false discovery rate (FDR) correction (*p.adjust* package). We considered a SNP significant if it exhibited an FDR < 0.2 and effect size > 0.5%. We used this approach to identify SNPs shifting systematically throughout the entire summer to fall transition (time points 1 to 11; before total population crash and reduction in mesocosm-wide replication of time point 12). We additionally used this approach to determine the interval length for which we had sufficient power to identify significant, relatively large effect size SNPs (FDR < 0.05 and >2 %) across all chromosomal arms (Fig. S4). Finally, we explored how SNP discovery changed when using intermediate time point data in the model. For example, for regressions of allele frequencies from t1->3, we compared the number of SNPs identified using data from t1, t2, and t3, as well only end time point data (i.e., just data from t1 and t3) (results in Fig. S4). The increase in SNP discovery using intermediate time point data informed our decision to incorporate intermediate time points in all regressions.

We used our GLM results to explore the genetic architecture of adaptive tracking. Specifically, we implemented a method developed and described by Rudman et al. 2022 to identify clusters of unlinked loci enriched in significant SNPs. Briefly, this method takes in scores each SNP as follows: 0: FDR > 0.2; 1: FDR < 0.2; 2: FDR < 0.05 and effect size > 2%; 3: FDR < 0.01 and effect size > 2%. The average SNP score is then calculated for sliding windows of 500 SNPs, using a 100 SNP step size. An empirical FDR for each window is generated by comparing observed scores to those generated for permuted windows, whereby SNP site information is randomly shuffled and window scores are re-computed. Windows with an empirical FDR < 0.05 are defined as significantly enriched and overlapping enriched windows are merged into a single, enriched windows. To winnow the list of enriched windows to a set of putatively unlinked (independently segregating windows) for each comparison, the squared correlation coefficient of founder genotypes is calculated between all pairs of significant SNPs within 1 Mb of each other. If the average SNP-pair linkage between two clusters is > 0.03, the clusters are merged. This process proceeds iteratively until all pairs of clusters exhibited average linkage < 0.03 (source code provided at https://github.com/greensii/dros-adaptive-tracking).

### Quantifying fluctuating selection using leave-one-out cross validation

We quantified the degree of parallel, neutral, and/or anti-parallel evolutionary change for individual replicate cages by re-implementing our GLM analysis using a leave-one-out cross-validation approach: sets of GLM significant SNPs were identified iteratively in *n* –1 ‘training’ cages, and the frequency dynamics of the rising alleles (FDR < 0.2, allele frequency change > 0.5 %) were then quantified in the left-out, ‘test’, cage. A set of control SNPs were selected for every set of target SNPs, whereby each control SNP was matched on the following: chromosomal arm, +/− 5% baseline population frequency, =/-0.5 cM/MB recombination rate, and inversion status (i.e., inside or outside of an inversion). The distributions of shifts between sets of target and matched control SNPs were then compared using a paired t-test to: 1) determine whether the magnitude of target shifts significantly exceeded (FDR < 0.05) background/neutral allele frequency change and 2) infer whether the direction of non-neutral shifts were in a predominantly concordant/parallel direction as that observed in the training cages (positive median allele frequency shift), or exhibited significantly anti-parallel behavior (negative median allele frequency shift). For a given left-out cage and set of target SNPs, changes between parallel and anti-parallel behavior across test intervals was indicative of a fluctuation in the dominant direction of selection. As target SNP behavior was always measured in the left-out cage, inference of fluctuating selection was insensitive to statistical artifacts, such as regression to the mean.

We first implemented this GLM and leave-one-out approach using sets of target SNPs generated across the entire summer to fall transition (t1->11). The frequency dynamics of target SNPs within left-out cages were measured from t1->11 (the interval over which they were identified in the training cages), separately for each chromosomal arm and genome-wide. Next, to resolve the pace of adaptive tracking in the system and explore the scale at which signals of fluctuating selection arise, we re-measured allele frequency shifts of t1->11 SNPs in increasingly shorter intervals. Specifically, we measured SNP behavior in all intervals starting at t1 and of decreasing length from t1->10 (11 weeks) through t1->2 (1 week). The sustained deterministic behavior of SNPs across replicates during the t1->2 interval motivated assessment of SNP behavior in each pair of single time points (t1->2, t2->3…t11->12), the first nine of which are single week intervals. This final pair of single time points (t11->12) includes the first observed freezing of the year (Fig. 1), and the total crash of the adult fly population within each replicate.

We next aimed to quantify the pervasiveness of fluctuating selection throughout the experimental period. Accordingly, we re-conducted our GLM and leave-one-out approach in all interval lengths greater than five time points (e.g., t1-6…t7-12) (a five time point interval length represented the shortest producing substantial, genome-wide GLM signal across all chromosomal arms; Fig. S4). For each set of test SNPs, we measured their allele frequency shifts across the time points over which they were identified, as well as all single time point comparisons (e.g., for SNPs identified during t1->6, we quantified allele frequency shifts from t1->6, as well as t1->2, t2->3…t11->12). For each set of target SNPs, we excluded sites identified as strongly parallel (FDR < 0.05) throughout the summer to fall transition (t1->11). This approach thus explicitly eliminates those sites most predominantly subject to primarily directional selection throughout the experimental period to explore whether finer modalities of environmental change drive systematic shifts in allele frequencies across cages. Finally, we quantified the changes in the dominant direction of selection throughout the experiment for each individual cage by enumerating the number of instances sets of target SNPs switched from being predominantly favored (significantly positive median phased allele frequency change) to predominantly selected against (significantly positive median phased allele frequency change), or vice versa, across all pairs of single time points.

### Quantifying the predictability of independent bouts of adaptive tracking

We leveraged results reported in Rudman et al. (2022) to probe the extent to which independent bouts of adaptive tracking in the system were underpinned by predictable shifts in genomic variation. The Rudman et al. study took place during the 2014 spring to fall transition, using the same outdoor mesocosm system (with a total of 10 cage replicates cages), and a total of 81 inbred strains, 51 of which were also used in this study (the 25 strains unique to this study were derived from flies collected at the same local Pennsylvania orchard population as overlapping strains). Rudman et al. collected samples for analysis of genomic variation monthly (five total time points, spanning 25 July to 7 November) and used identical methods of haplotype-derived allele frequency estimation, GLM-based identification of sites moving systematically through time, and clustering of enriched regions of genomic variation.

We first used PCA to explore the impact of experimental year and collection time on variance in allele frequencies across all samples from studies. While the experiments were initiated on different calendar days and extended different periods of time, we standardized collection time by days since initial founding of the mesocosm replicates. We projected each sample onto the first two principal components to ask – are variance in allele frequencies predominantly driven by experimental year and/or collection time? Next, we tested whether, despite inter-annual differences in founder population allele frequencies and environmental conditions, a parallel set of SNPs underpinned seasonal adaptation across years. We compared the allele frequency shifts obtained across all interval lengths and comparisons of this study (e.g., all two time point to eleven time point intervals), to each of the four monthly intervals of the Rudman et al. study (270 total comparisons). For each comparison, we computed Pearson correlations between observed allele frequency shifts (∼1.6 M SNPs with overlapping data across studies). Positive correlations indicated that SNPs moved in a concordant manner between the tested 2014 and 2021 intervals, and negative correlations to SNPs moving in predominantly opposing directions between years. To infer whether observed correlations exceeded those expected by chance, we computed a distribution of null correlations by randomly shuffling the allele frequency shift values by SNP for each comparison (SNPs shuffled to sites matched on chromosomal arm, starting frequency, recombination rate, and inversion status). We compared the distribution of observed and shuffled correlations using an F test to determine whether an excess of parallel (and anti-parallel) intervals were observed between years. We re-conducted this analysis separately for each chromosomal arm. Finally, to explore whether those intervals with heightened allele frequency concordance across years aligned with conspicuous shifts in the selective environment, we identified the interval of the current study that was best matched (via highest genome-wide correlation) to each of the four monthly intervals of the 2014 experiment.

## Acknowledgements

We are grateful to members of the Petrov and Schmidt labs for discussion during experimental design, lab work, and data analysis. We acknowledge Marianthi Karageorgi and Katie Solari for support with sample sequencing. We thank Jess Rhodes and Sharon Greenblum for feedback on statistical analysis and computing. We additionally thank Moi Exposito-Alonso, Hunter Fraser, Olivia Ghosh, Sharon Greenblum, James Hemker, Marianthi Karageorgi, Bernard Kim, Jess Rhodes, Katie Solari, and Sophie Walton for insightful comments on drafts of this manuscript.

## Author Contributions

The experimental was conceived by M.C.B., D.A.P, and P.S. The mesocosm study was conducted by M.C.B., S.B., H.O., and P.S. Population census size estimates were conducted by A.H. DNA extraction, library preparation, and sequencing was conducted by M.C.B. Bioinformatic analysis and statistical analysis of allele frequency data was carried out by M.C.B and D.A.P. M.C.B., D.A.P, and P.S. wrote the manuscript, with feedback from all authors.

## Materials & Correspondence

Correspondence and materials requests should be submitted to Mark Bitter, Paul Schmidt, and Dmitri Petrov.

## Supplementary Material

### Supplementary Figures

**Figure S1.**
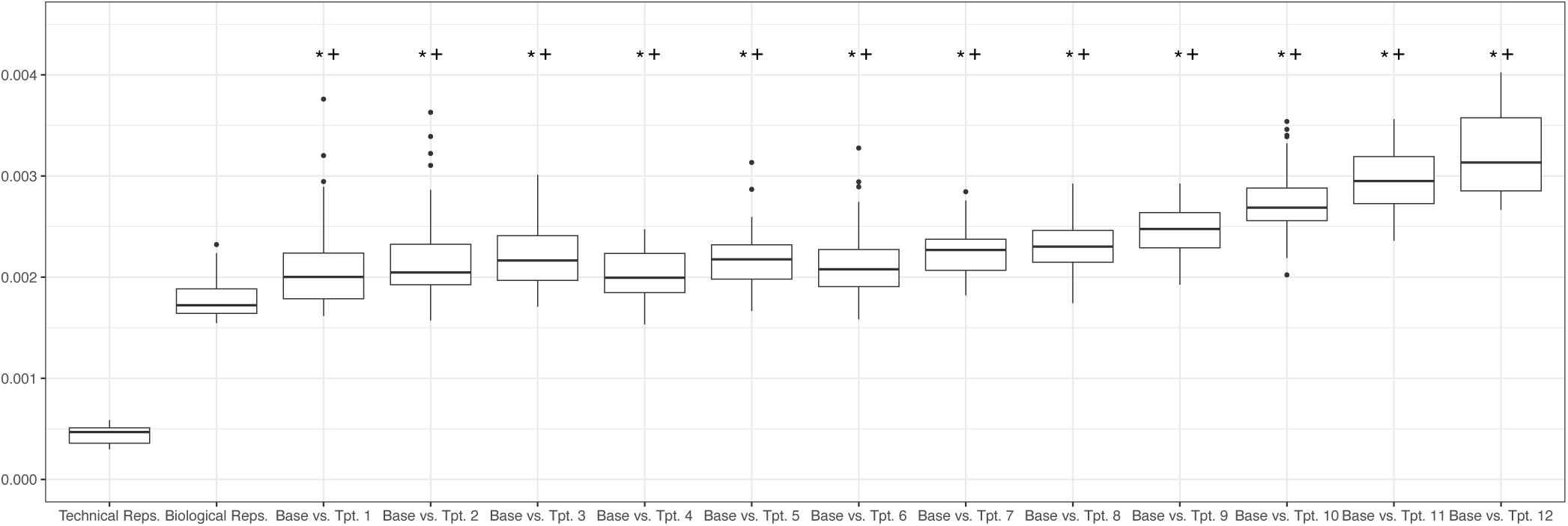
Genome-wide differentiation throughout experiment. | Distribution of genome-wide Fst computed pairwise between all technical replicate samples (same DNA extraction, different library preparation and sequencing lane; N = 10), biological replicates (same time point and replicate cage, different pool of 100 flies; N = 17), and samples collected at each time point (N = 12) and the baseline population samples (‘Base’, N = 4), for each of the twelve time points. The distributions of genome-wide F_ST_ between time point samples and the baseline population was significantly greater than that observed between technical replicates (*) and biological replicates (+) (paired T test; P-value < 0.01). F_ST_ between time point samples and the baseline population increased throughout the course of the experiment (generalized linear model; P = 0.039).

**Figure S2.**
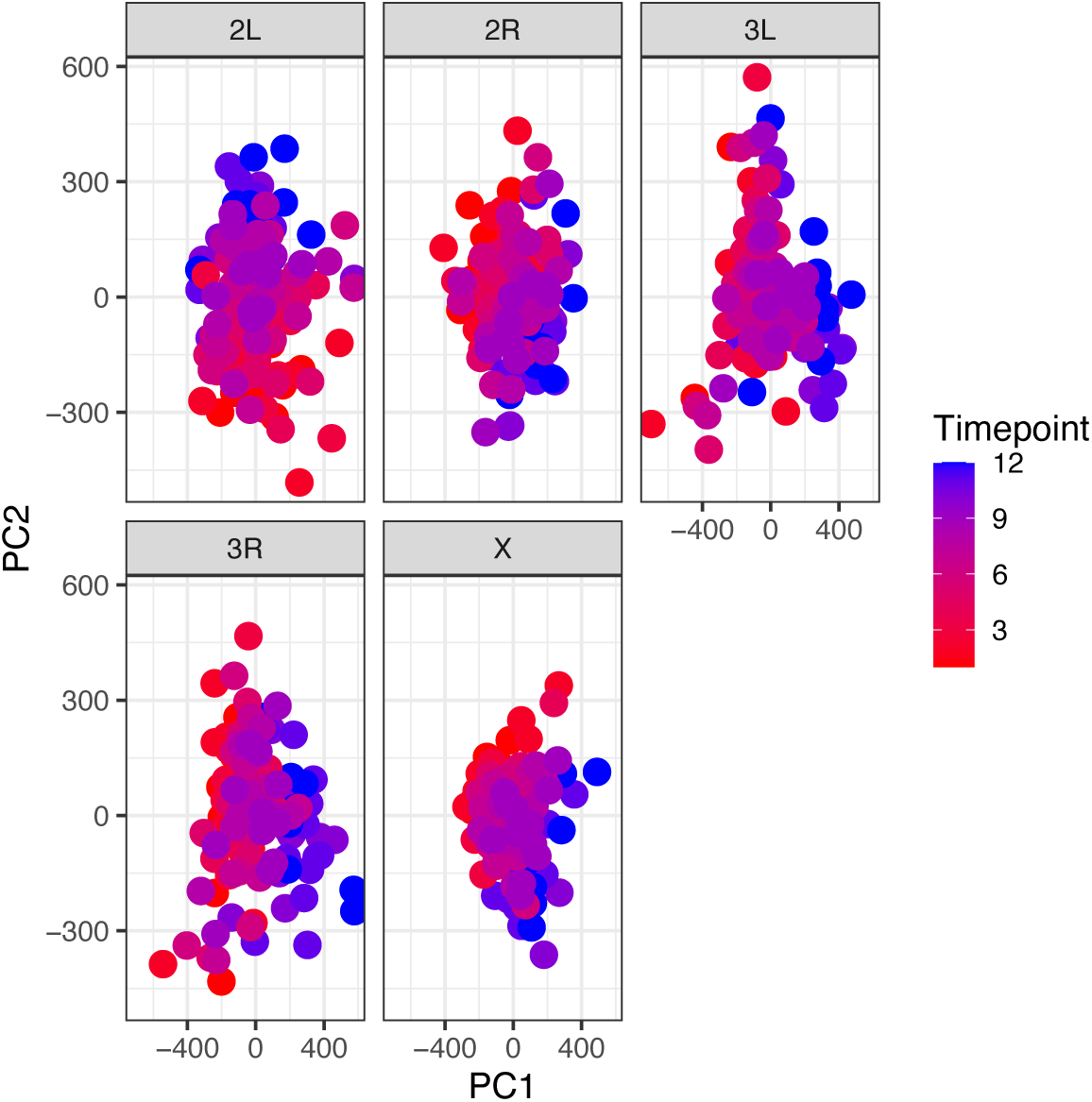
Principal component analysis by chromosomal arm. | Principal component analysis of normalized allele frequency data across all replicates and time points, run separately on SNPs of each chromosomal arm (2L: 453,445 SNPs; 2R: 373,755 SNPs; 3L: 445,457 SNPs; 3R: 448,008 SNPs; X: 254,497. Sample scores are projected onto the first two principal components and colored by collection time point. Variance explained for PC1 and PC2 for each chromosomal arm were as follows: 2L: 7.12% and 6.33%; 2R: 5.91% and 5.04%; 3L: 7.75% and 5.60%; 3R: 7.42% and 5.82%; X: 6.71% and 5.29%.

**Figure S3.**
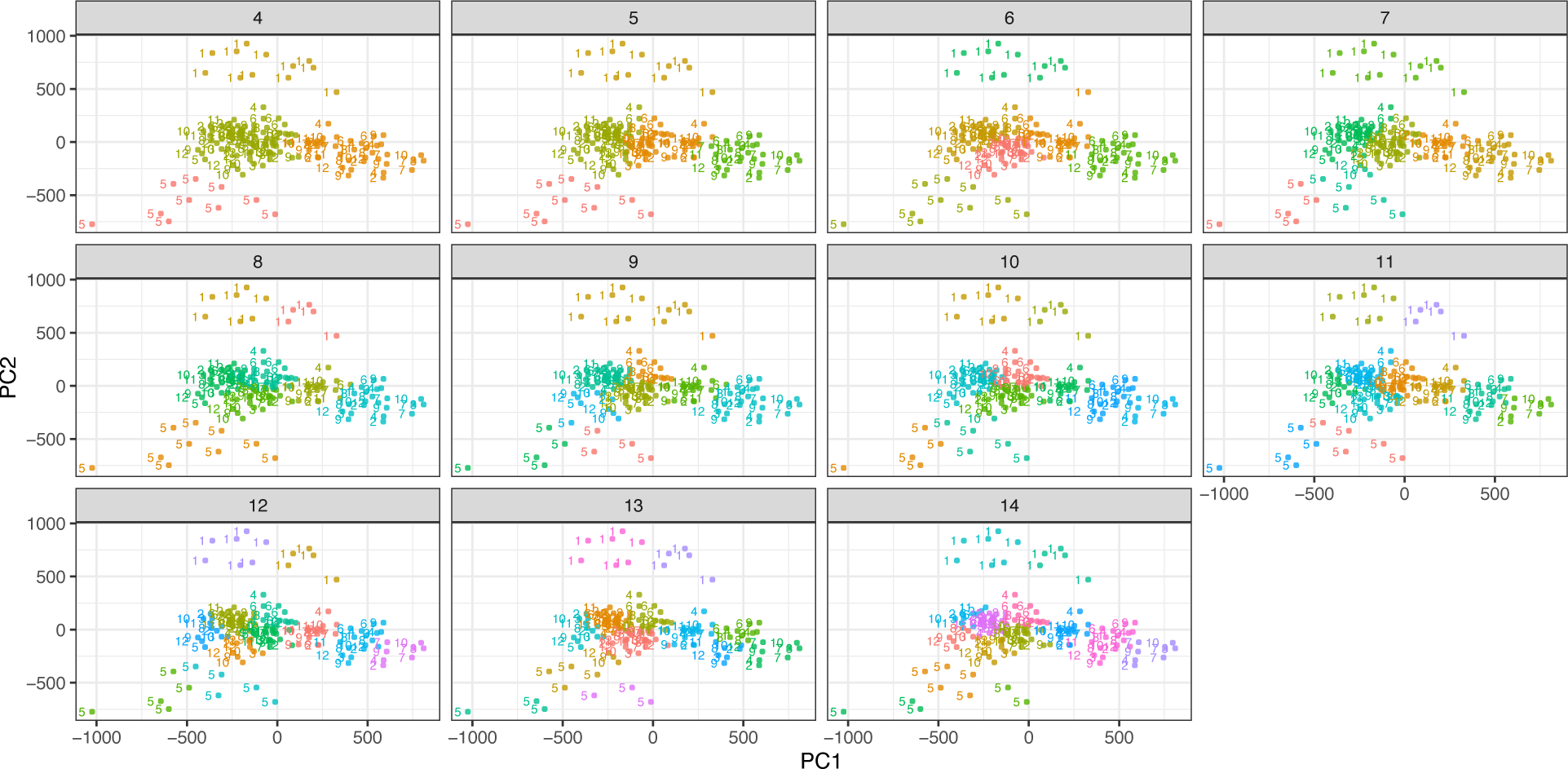
K-means clustering of PCA. | Unsupervised K-means clustering was conducted using each sample’s PC 1 and 2 scores (main text Figure 2A/B). Depicted here are results for K4-14 in which individual samples are colored by cluster membership and labelled by replicate cage.

**Fig. S4.**
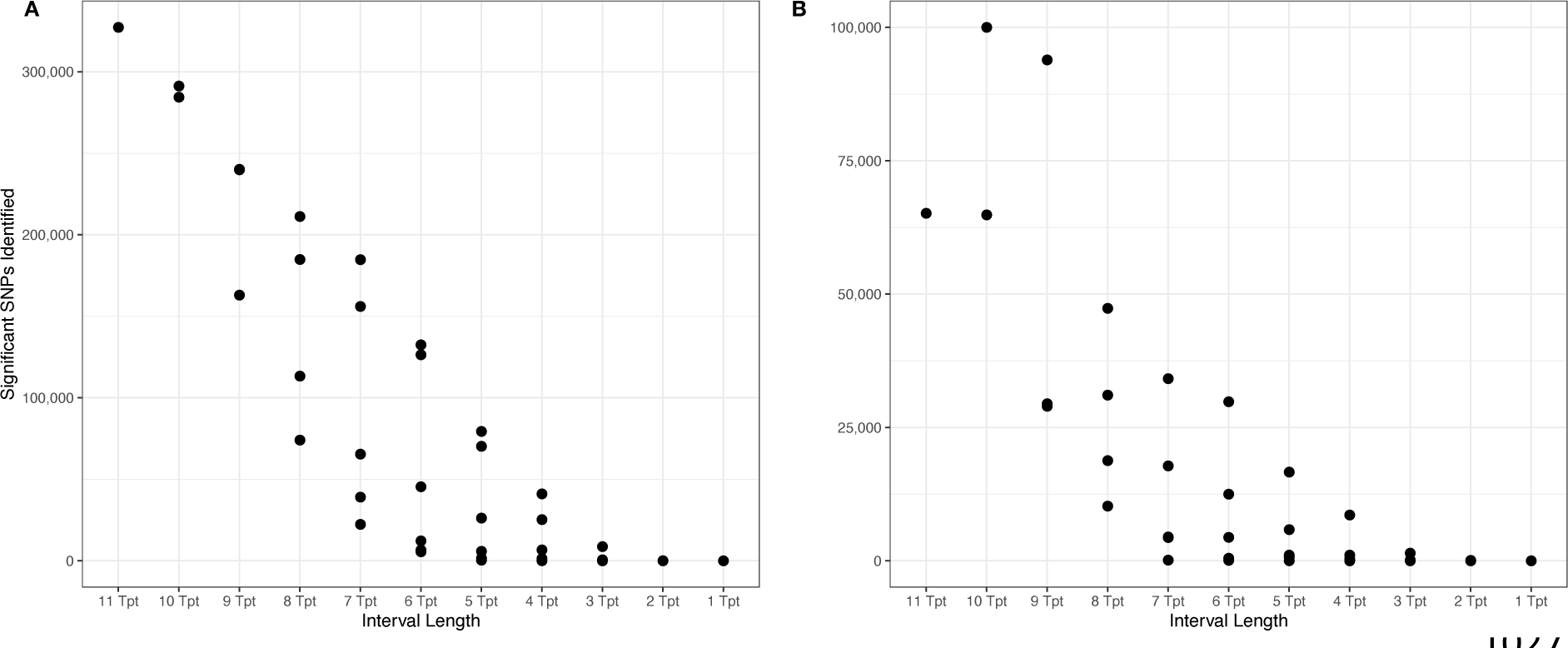
SNP discovery as a function of GLM interval length. | The number of SNPs with Benjamini-Hochberg FDR < 0.05 and effect size > 1% identified via regression of allele frequencies through time using all possible interval lengths. For each interval length, all possible time point comparisons were considered (each point represents a separate comparison). For example, nine time point intervals include regression of allele frequencies from t1->10, t2->11, and t3->12. (A) Depicts results for GLMs using intermediary time points within each comparison in the regression (e.g., for t1->3, model included allele frequency data from t1, t2, and t3). (B) Depicts results for GLMs in which only allele frequency data from the first and last time point of the comparison were considered in the model (e.g., for t1->3, model included allele frequency data from only t1 and t3).

**Figure S5.**
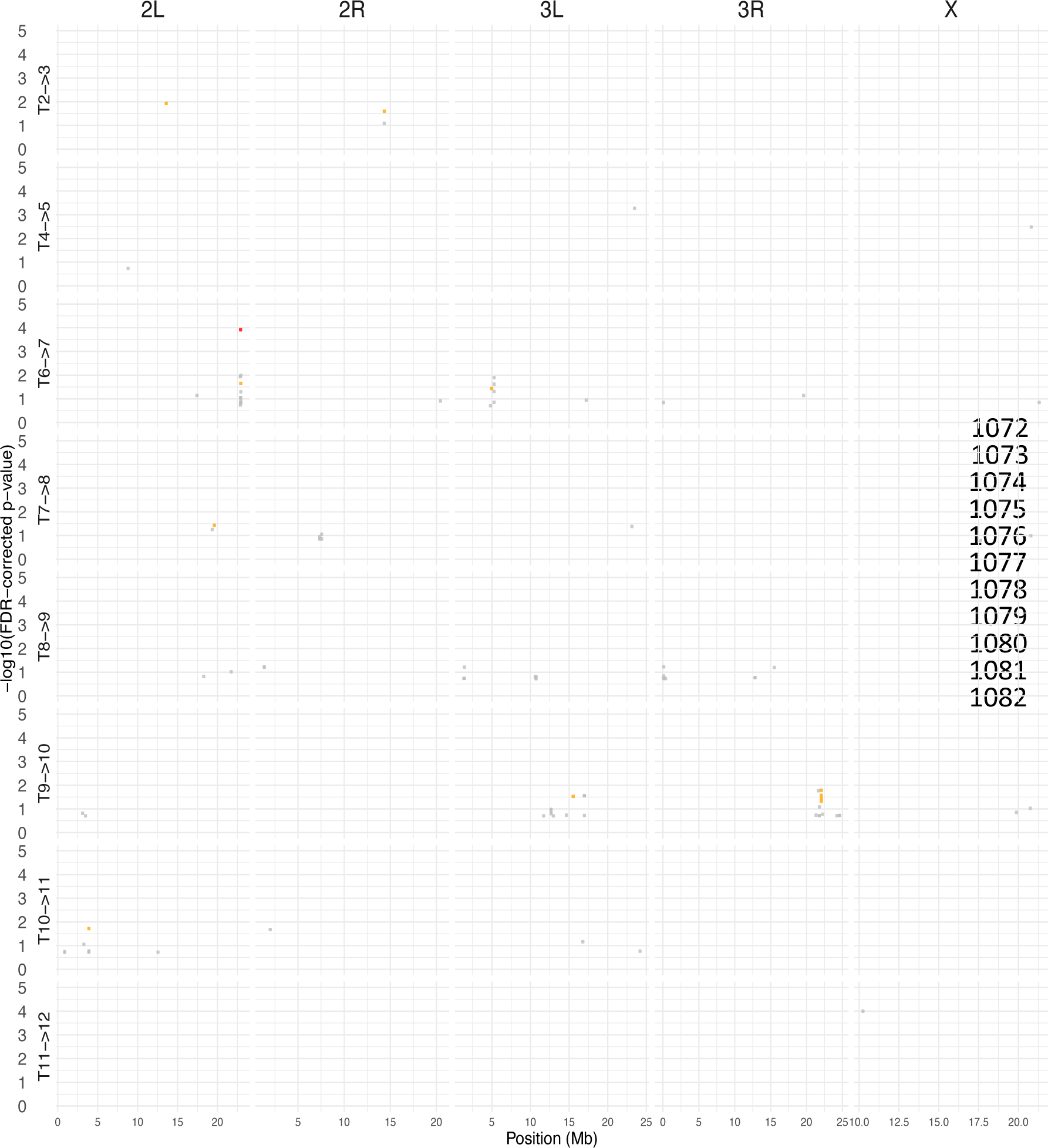
Genomic distribution of SNPs identified as significant using a generalized linear model and regression of allele frequencies across all single pairs of time points. | SNPs (points) are plotted as a function of chromosomal location and Benjamini-Hochberg FDR-corrected P value. SNPs are colored according to FDR-corrected P value and effect size: grey = FDR < 0.2, effect size > 0.05%; orange = FDR < 0.05 and effect size > 2%; red = FDR < 0.01 and effect size > 2%. Note t1->2 not shown as no significant SNPs were identified in that interval.

**Figure S6.**
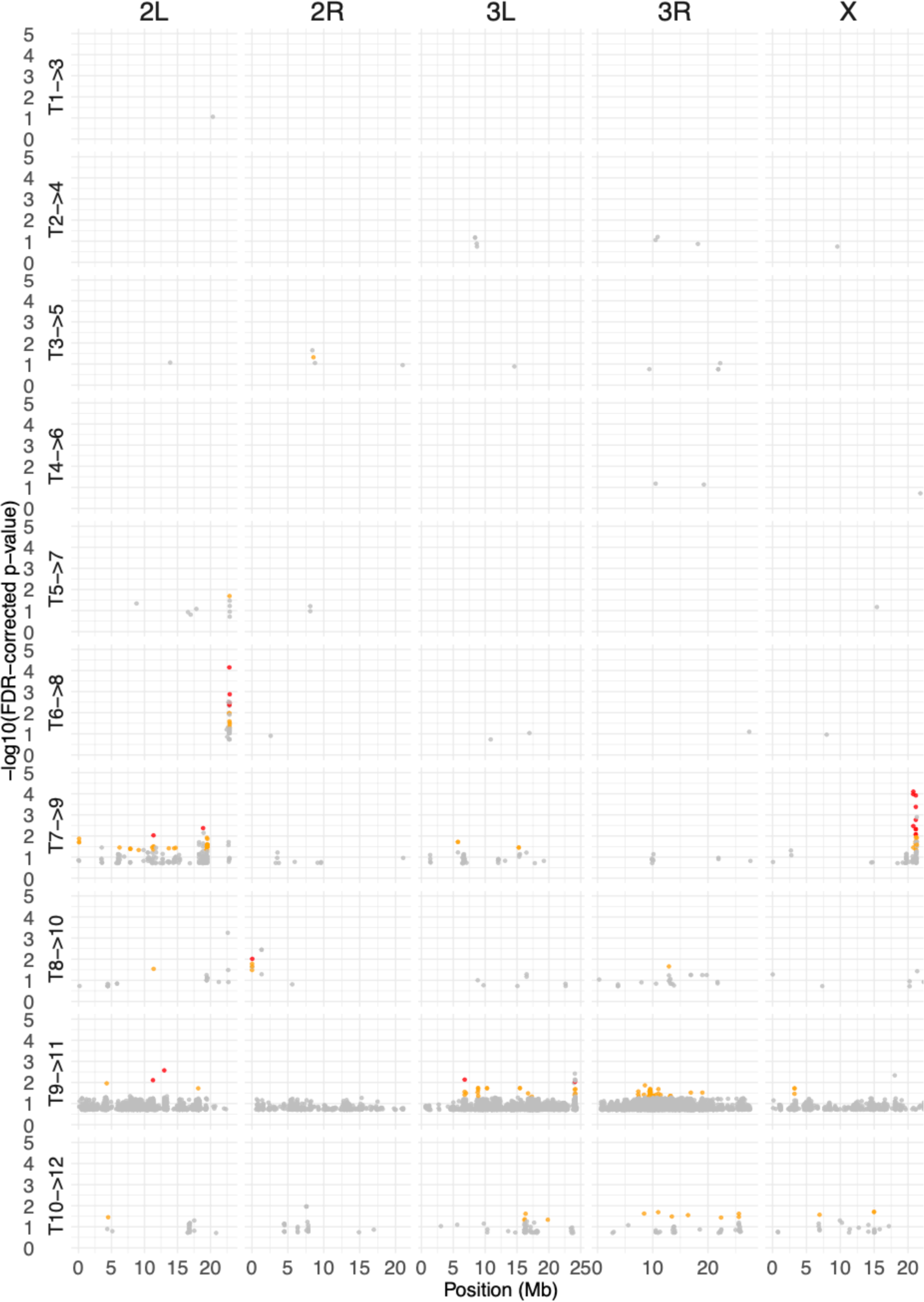
Genomic distribution of SNPs identified as significant using a generalized linear model and regression of allele frequencies across all two time point intervals. | SNPs (points) are plotted as a function of chromosomal location and Benjamini-Hochberg FDR-corrected P value. SNPs are colored according to FDR-corrected P value and effect size: grey = FDR < 0.2, effect size > 0.05%; orange = FDR < 0.05 and effect size > 2%; red = FDR < 0.01 and effect size > 2%.

**Figure S7.**
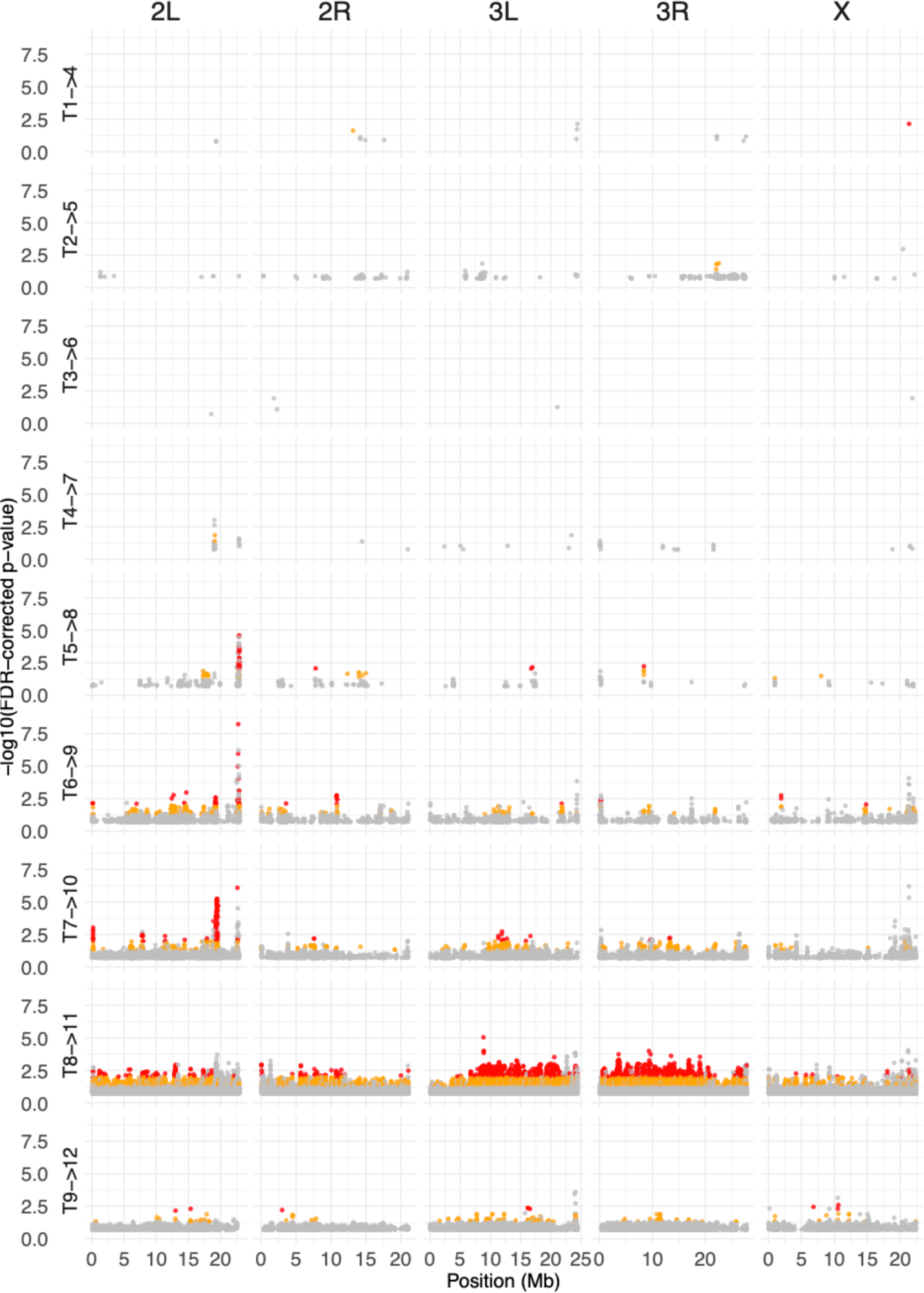
Genomic distribution of SNPs identified as significant using a generalized linear model and regression of allele frequencies across all three time point intervals. | SNPs (points) are plotted as a function of chromosomal location and Benjamini-Hochberg FDR-corrected P value. SNPs are colored according to FDR-corrected P value and effect size: grey = FDR < 0.2, effect size > 0.05%; orange = FDR < 0.05 and effect size > 2%; red = FDR < 0.01 and effect size > 2%.

**Figure S8.**
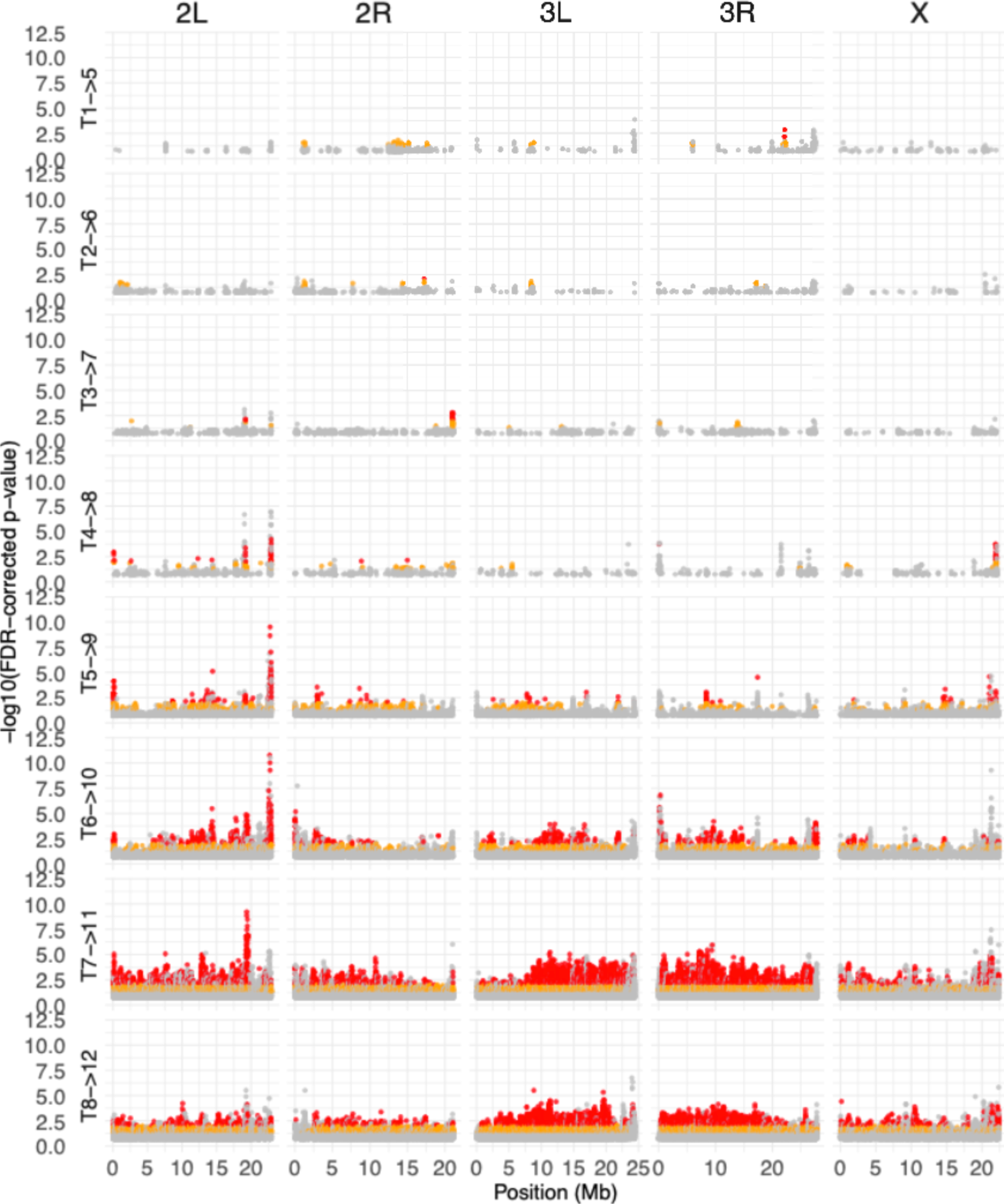
Genomic distribution of SNPs identified as significant using a generalized linear model and regression of allele frequencies across all four time point intervals. | SNPs (points) are plotted as a function of chromosomal location and Benjamini-Hochberg FDR-corrected P value. SNPs are colored according to FDR-corrected P value and effect size: grey = FDR < 0.2, effect size > 0.05%; orange = FDR < 0.05 and effect size > 2%; red = FDR < 0.01 and effect size > 2%.

**Figure S9.**
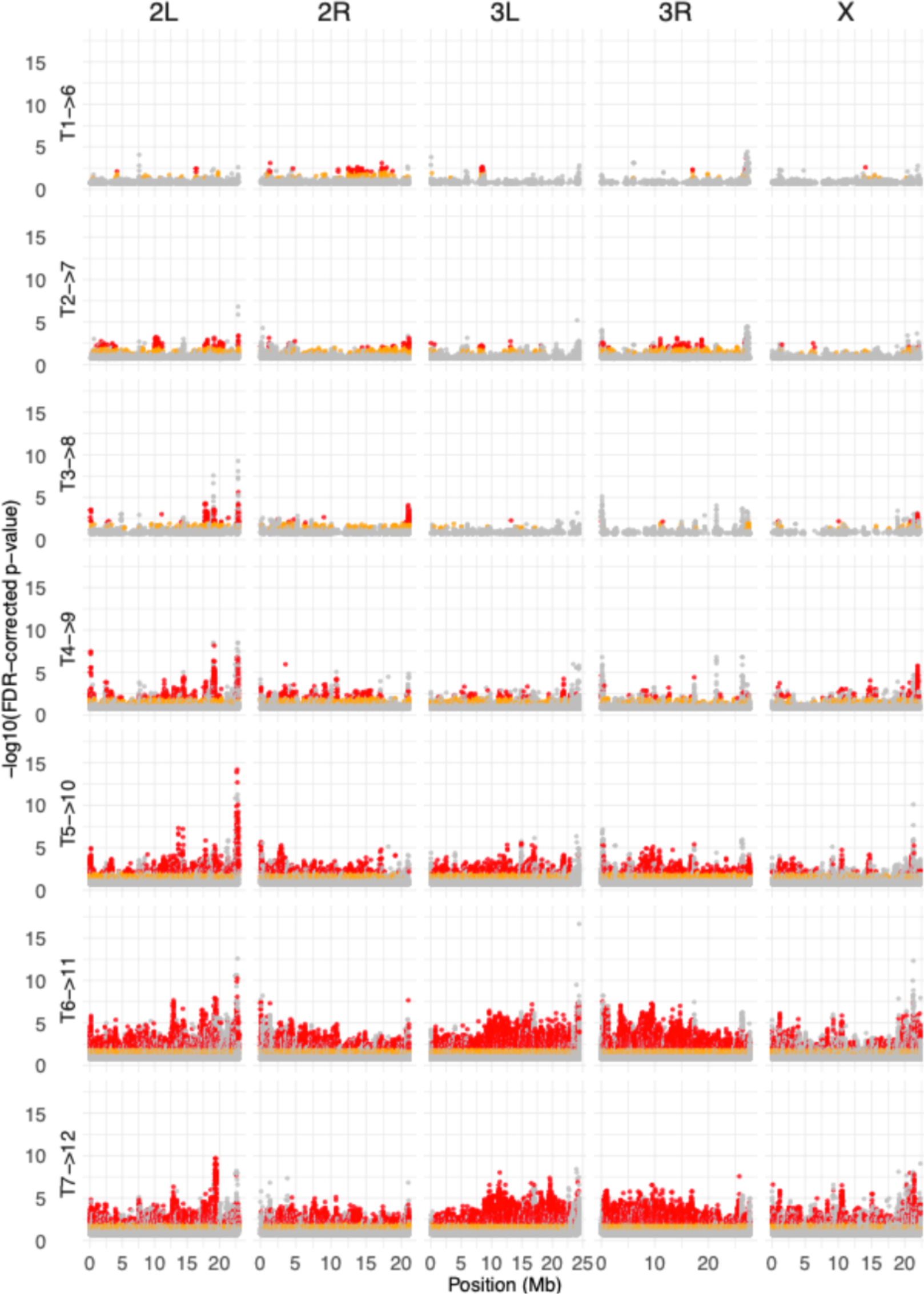
Genomic distribution of SNPs identified as significant using a generalized linear model and regression of allele frequencies across all five time point intervals. | SNPs (points) are plotted as a function of chromosomal location and Benjamini-Hochberg FDR-corrected P value. SNPs are colored according to FDR-corrected P value and effect size: grey = FDR < 0.2, effect size > 0.05%; orange = FDR < 0.05 and effect size > 2%; red = FDR < 0.01 and effect size > 2%.

**Figure S10.**
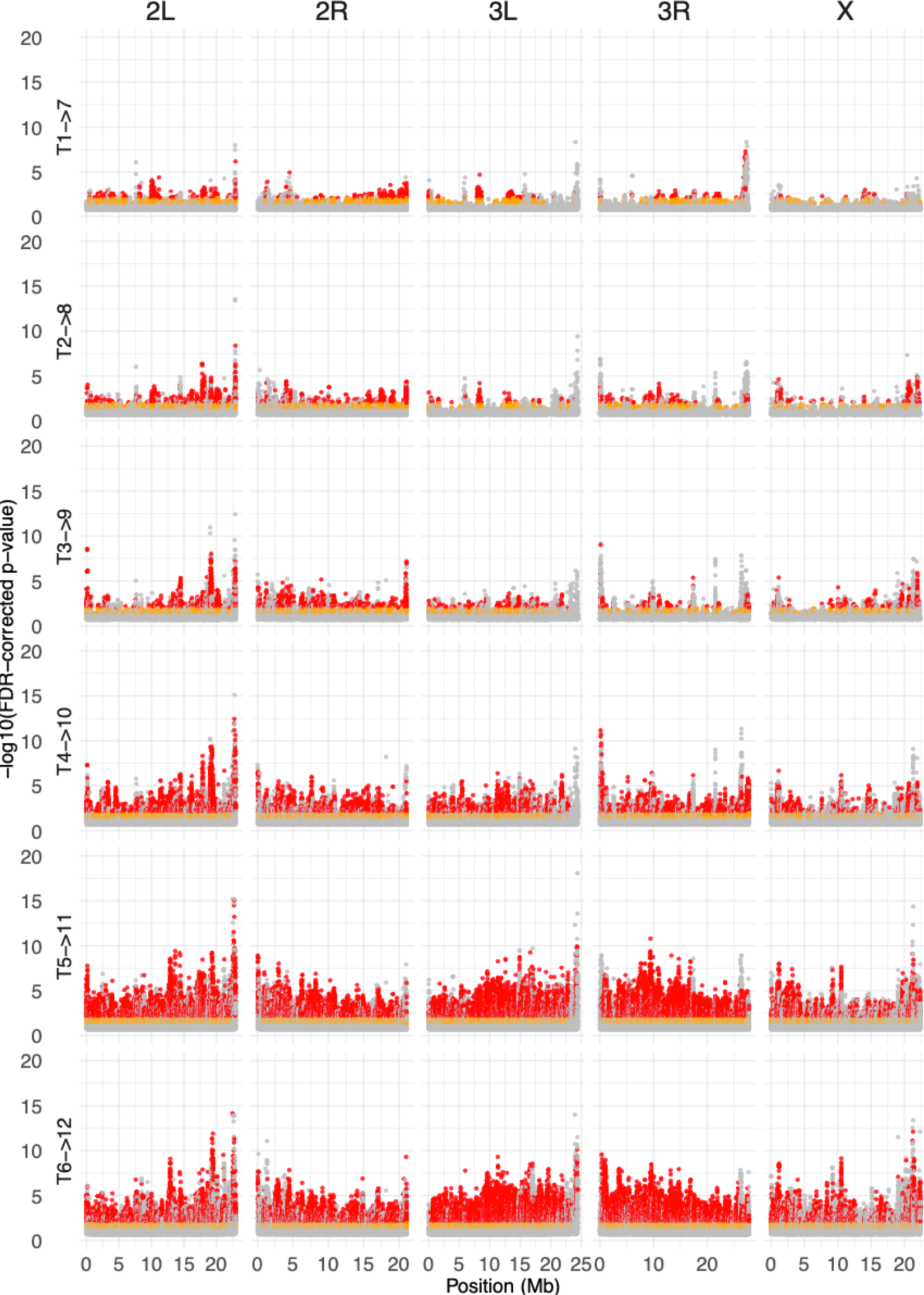
Genomic distribution of SNPs identified as significant using a generalized linear model and regression of allele frequencies across all six time point intervals. | SNPs (points) are plotted as a function of chromosomal location and Benjamini-Hochberg FDR-corrected P value. SNPs are colored according to FDR-corrected P value and effect size: grey = FDR < 0.2, effect size > 0.05%; orange = FDR < 0.05 and effect size > 2%; red = FDR < 0.01 and effect size > 2%.

**Figure S11.**
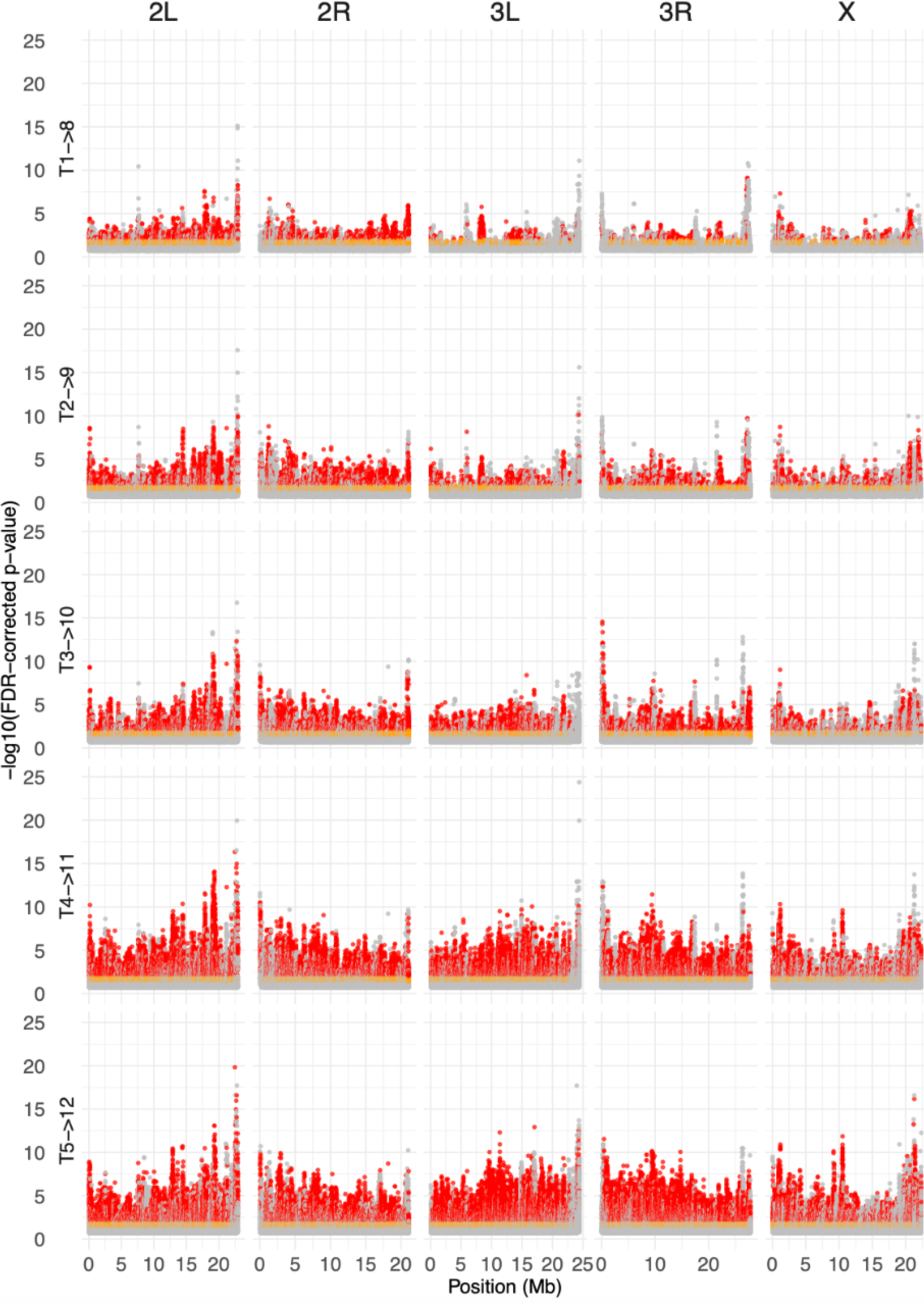
Genomic distribution of SNPs identified as significant using a generalized linear model and regression of allele frequencies across all seven time point intervals. | SNPs (points) are plotted as a function of chromosomal location and Benjamini-Hochberg FDR-corrected P value. SNPs are colored according to FDR-corrected P value and effect size: grey = FDR < 0.2, effect size > 0.05%; orange = FDR < 0.05 and effect size > 2%; red = FDR < 0.01 and effect size > 2%.

**Figure S12.**
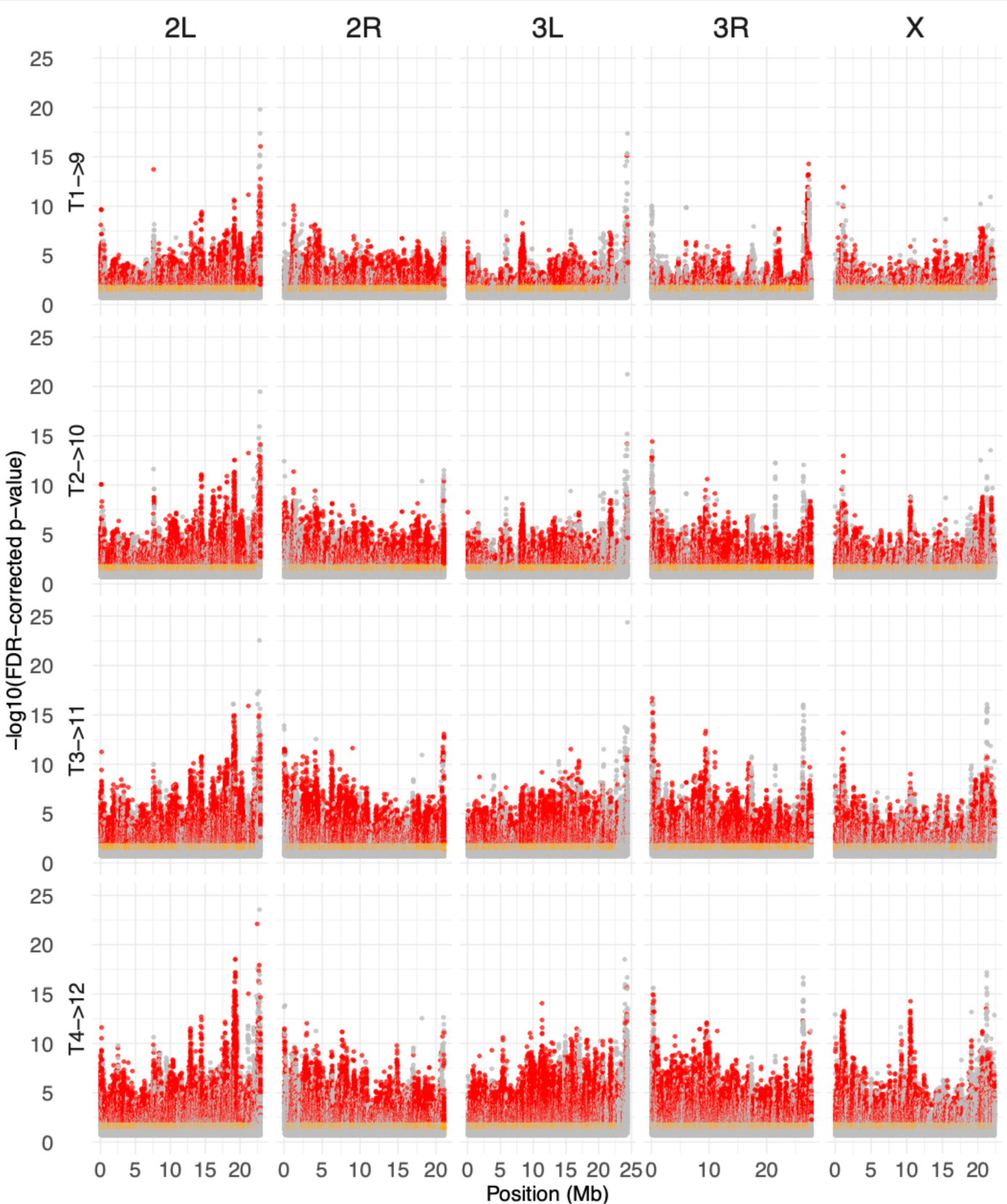
Genomic distribution of SNPs identified as significant using a generalized linear model and regression of allele frequencies across all eight time point intervals. | SNPs (points) are plotted as a function of chromosomal location and Benjamini-Hochberg FDR-corrected P value. SNPs are colored according to FDR-corrected P value and effect size: grey = FDR < 0.2, effect size > 0.05%; orange = FDR < 0.05 and effect size > 2%; red = FDR < 0.01 and effect size > 2%.

**Figure S13.**
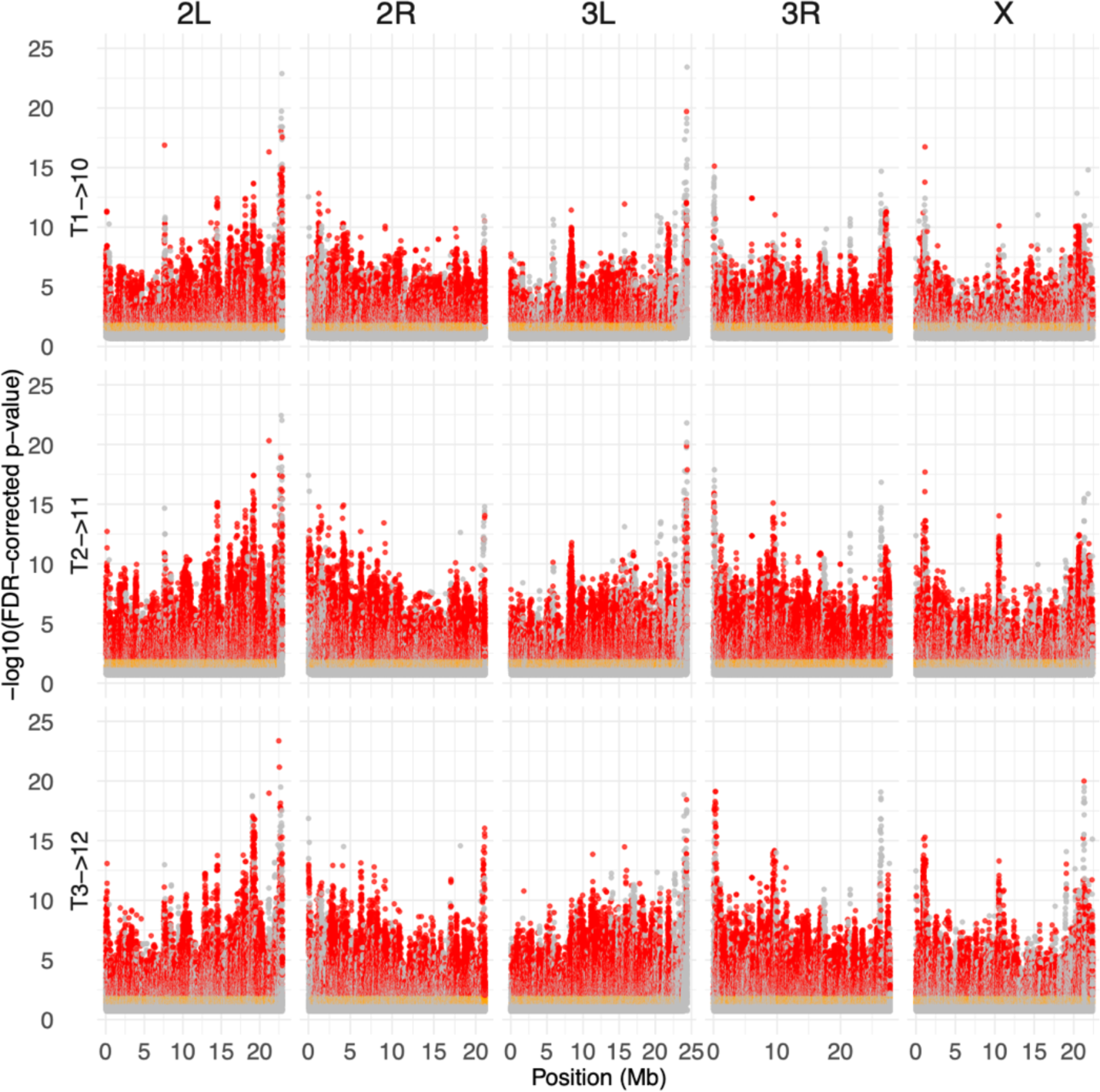
Genomic distribution of SNPs identified as significant using a generalized linear model and regression of allele frequencies across all nine time point intervals. | SNPs (points) are plotted as a function of chromosomal location and Benjamini-Hochberg FDR-corrected P value. SNPs are colored according to FDR-corrected P value and effect size: grey = FDR < 0.2, effect size > 0.05%; orange = FDR < 0.05 and effect size > 2%; red = FDR < 0.01 and effect size > 2%.

**Figure S14.**
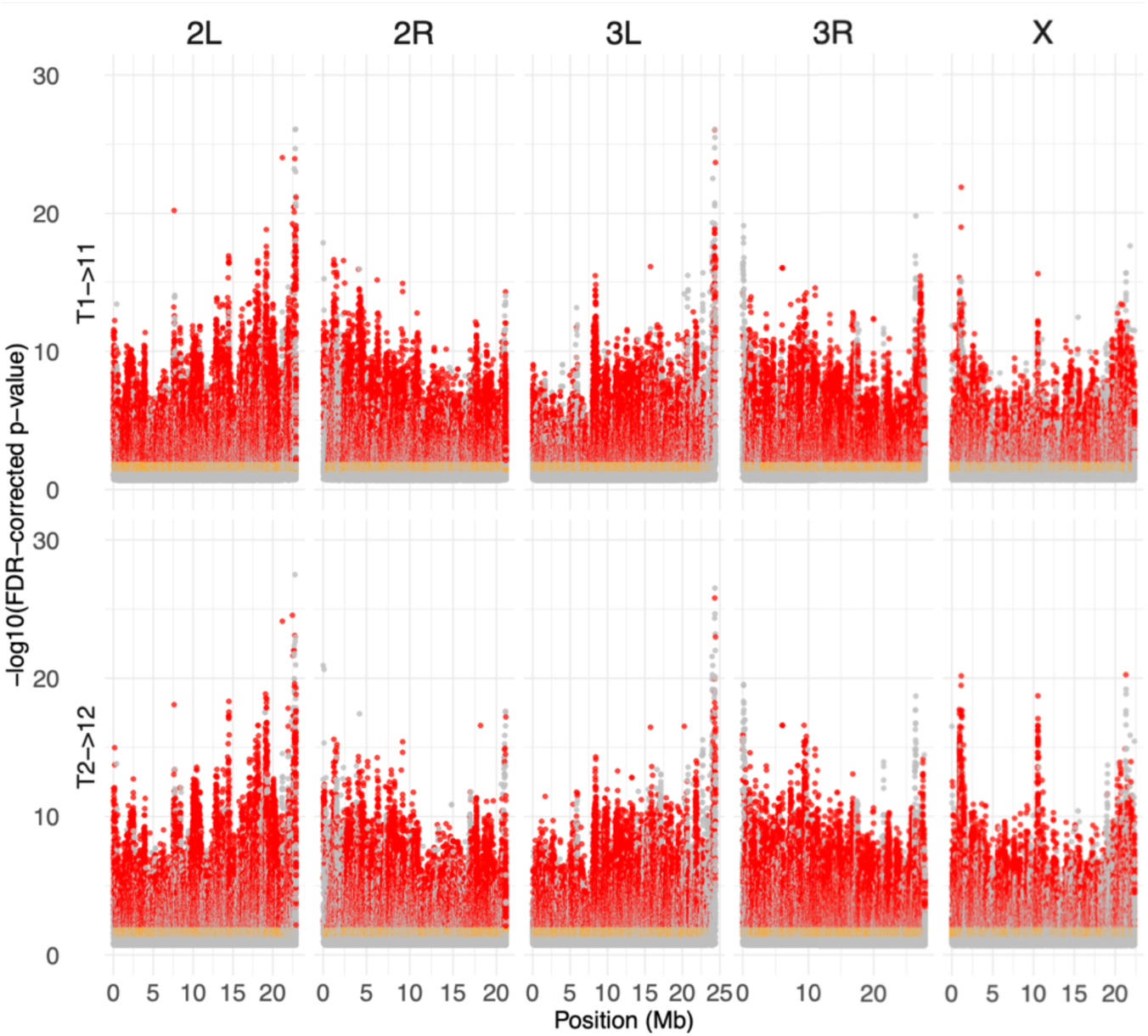
Genomic distribution of SNPs identified as significant using a generalized linear model and regression of allele frequencies across all ten time point intervals. | SNPs (points) are plotted as a function of chromosomal location and Benjamini-Hochberg FDR-corrected P value. SNPs are colored according to FDR-corrected P value and effect size: grey = FDR < 0.2, effect size > 0.05%; orange = FDR < 0.05 and effect size > 2%; red = FDR < 0.01 and effect size > 2%.

**Figure S15.**
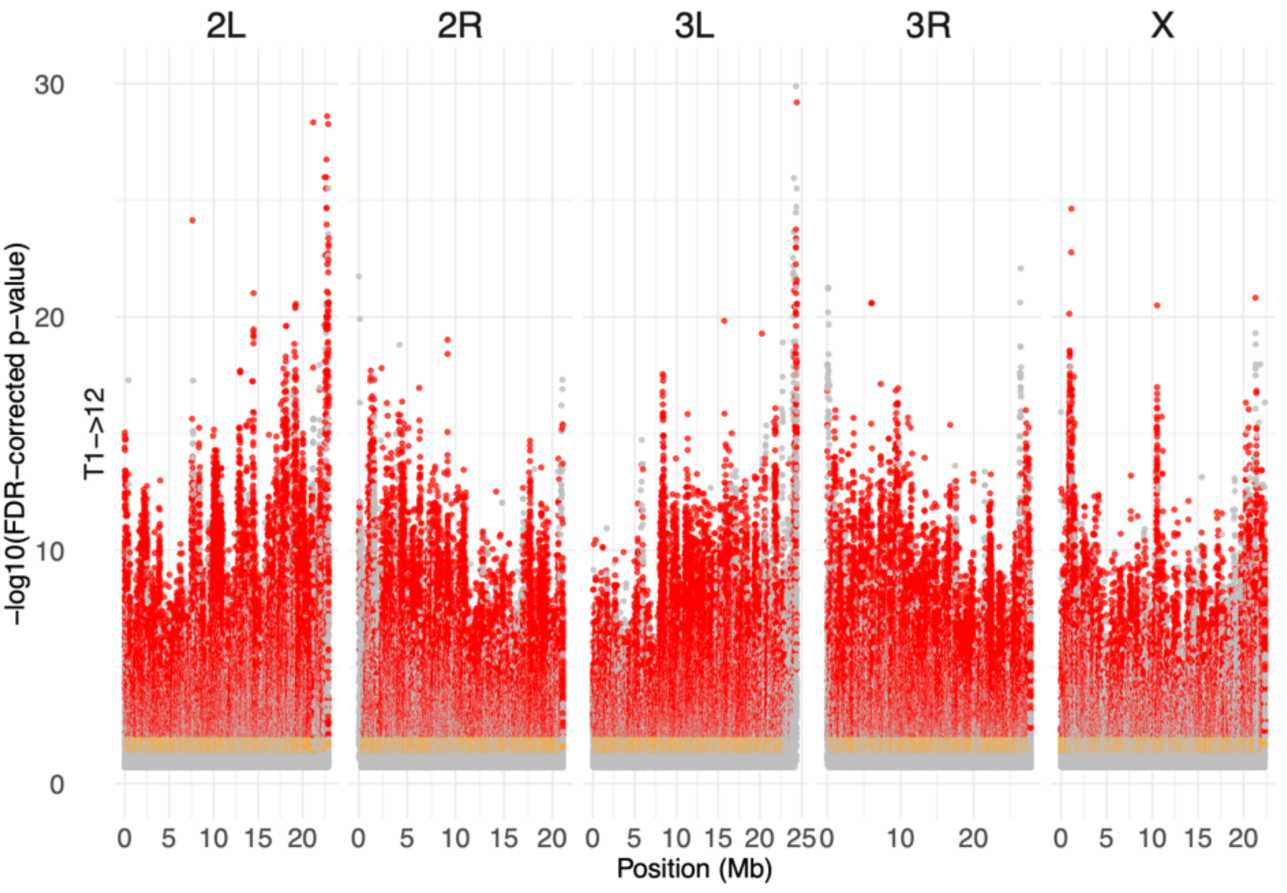
Genomic distribution of SNPs identified as significant using a generalized linear model and regression of allele frequencies across all ten time point intervals. | SNPs (points) are plotted as a function of chromosomal location and Benjamini-Hochberg FDR-corrected P value. SNPs are colored according to FDR-corrected P value and effect size: grey = FDR < 0.2, effect size > 0.05%; orange = FDR < 0.05 and effect size > 2%; red = FDR < 0.01 and effect size > 2%.

**Figure S15.**
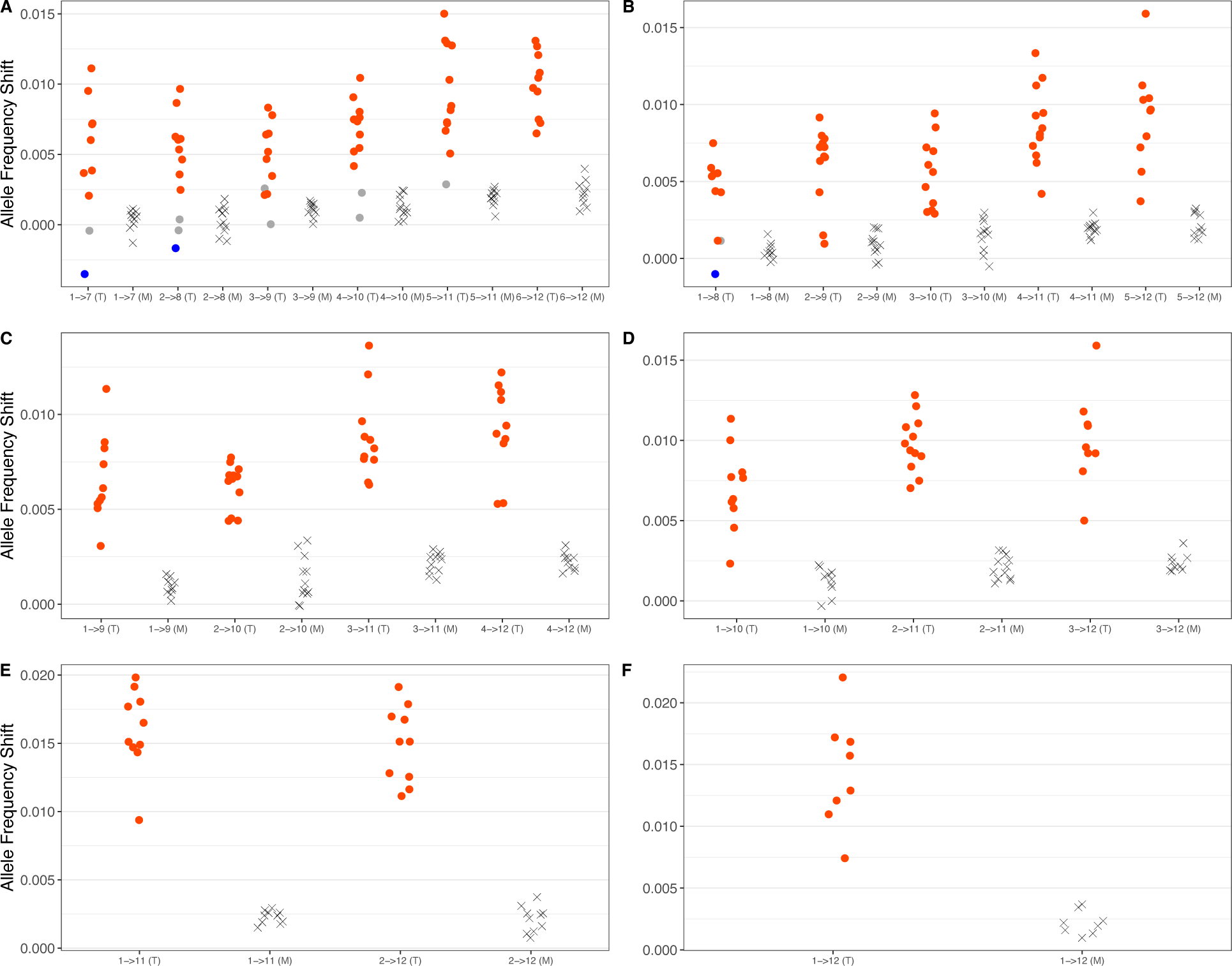
| Median allele frequency shifts for each replicate at target (T) SNPs (circles) and matched (M) controls (M, X’s) identified using a GLM and leave-one-out approach for all six (A), seven (B), eight (C), nine (D), ten (E), and eleven (F) time point intervals. Allele frequency shifts were measured, and are plotted, across the interval over which sets of test SNPs were identified. Median shifts at target sites are colored red or blue if the distribution of phased allele frequency shifts was significantly greater or less than that for matched control sites, respectively.

**Figure S16.**
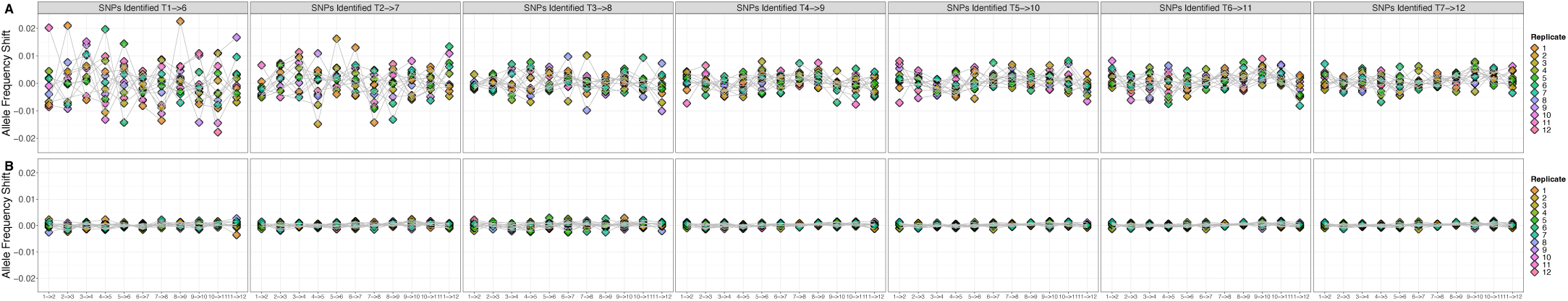
| Median shifts at target SNPs (A) and matched control SNPs (B) identified in each possible five time point interval and measured across each pair of single time points. Point color corresponds to replicate cage ID’s and grey lines connect the median shifts for each cage between consecutive time point intervals.

**Figure S17.**
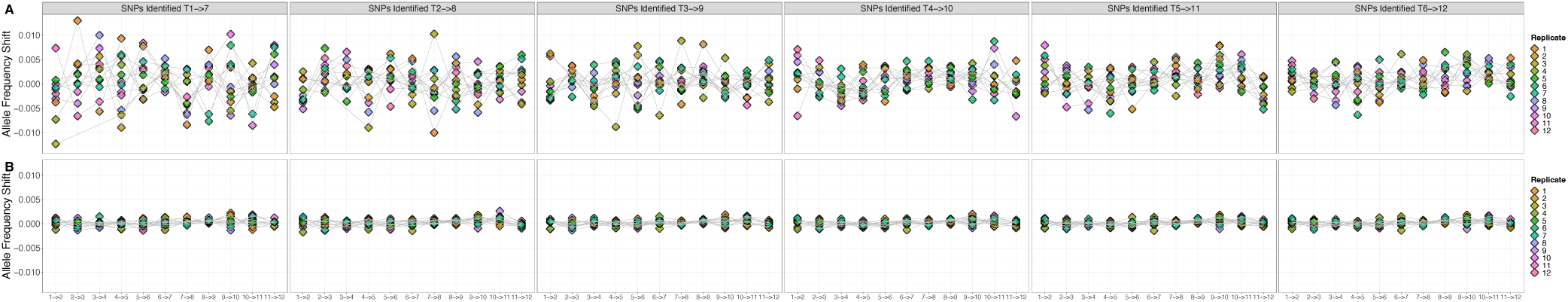
| Median shifts at target SNPs (A) and matched control SNPs (B) identified in each possible six time point interval and measured across each pair of single time points. Point color corresponds to replicate cage ID’s and grey lines connect the median shifts for each cage between consecutive time point intervals.

**Figure S18.**
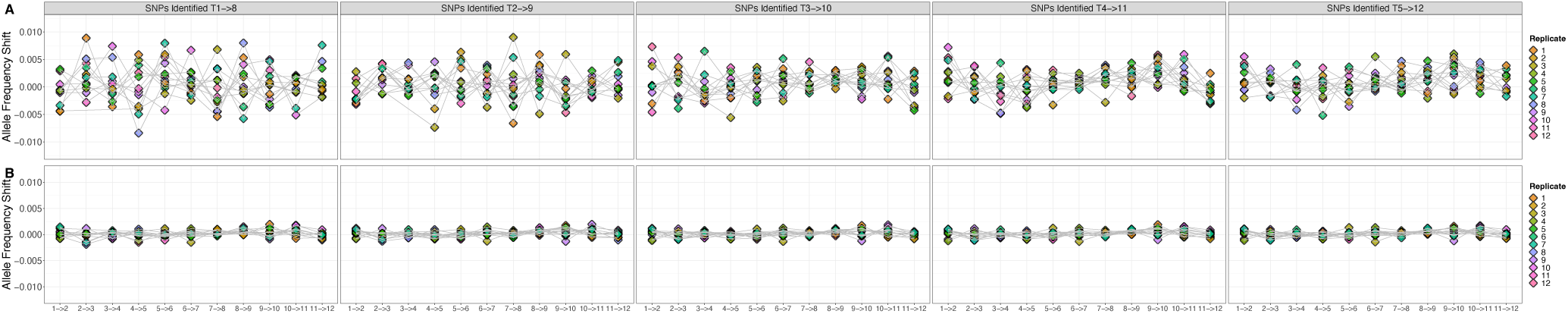
| Median shifts at target SNPs (A) and matched control SNPs (B) identified in each possible seven time point interval and measured across each pair of single time points. Point color corresponds to replicate cage ID’s and grey lines connect the median shifts for each cage between consecutive time point intervals.

**Figure S19.**
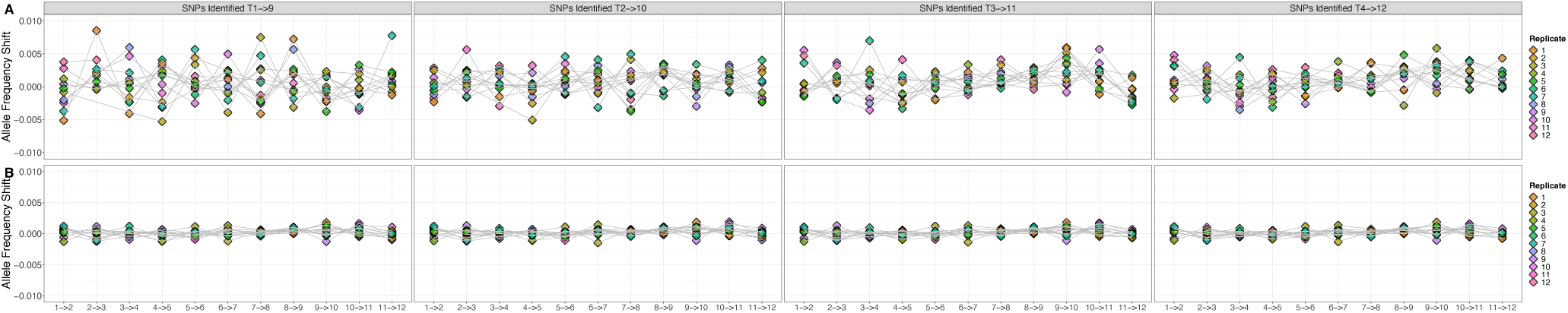
| Median shifts at target SNPs (A) and matched control SNPs (B) identified in each possible eight time point interval and measured across each pair of single time points. Point color corresponds to replicate cage ID’s and grey lines connect the median shifts for each cage between consecutive time point intervals.

**Figure S20.**
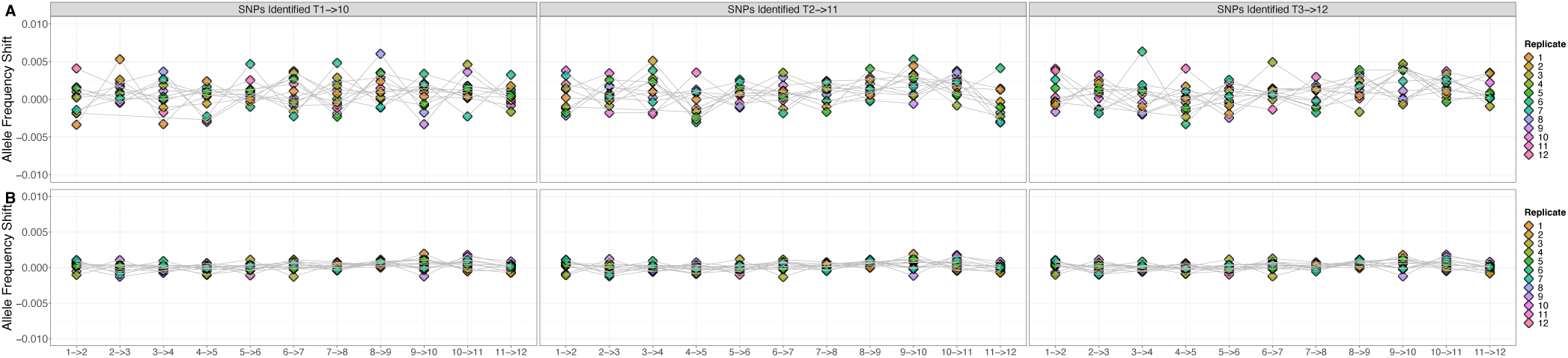
| Median shifts at target SNPs (A) and matched control SNPs (B) identified in each possible nine time point interval and measured across each pair of single time points. Point color corresponds to replicate cage ID’s and grey lines connect the median shifts for each cage between consecutive time point intervals.

**Figure S21.**
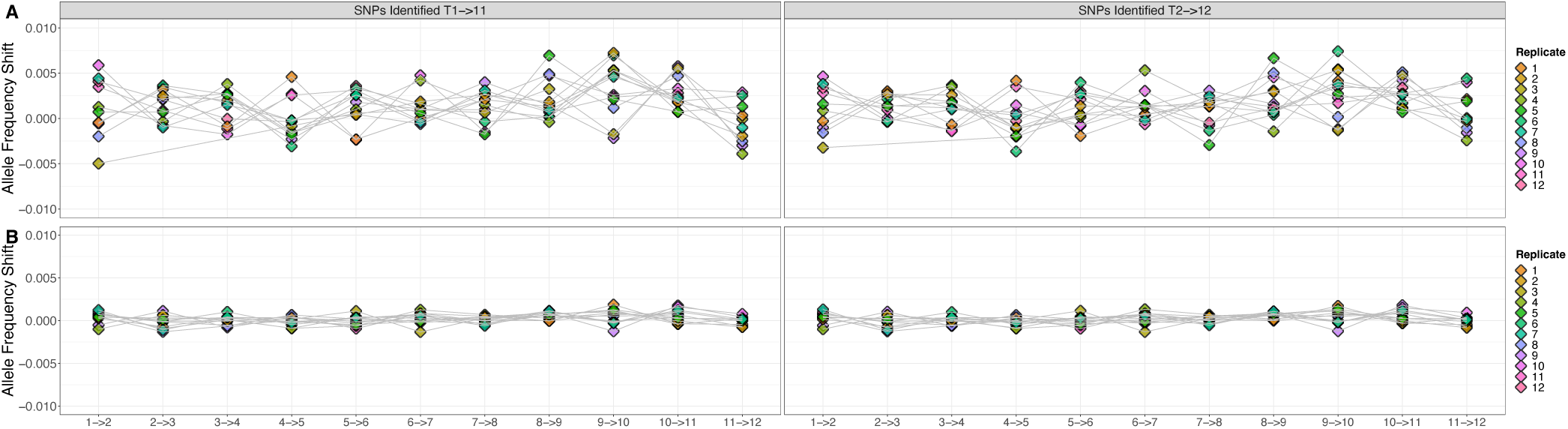
| Median shifts at target SNPs (A) and matched control SNPs (B) identified in each possible ten time point interval and measured across each pair of single time points. Point color corresponds to replicate cage ID’s and grey lines connect the median shifts for each cage between consecutive time point intervals.

**Figure S22.**
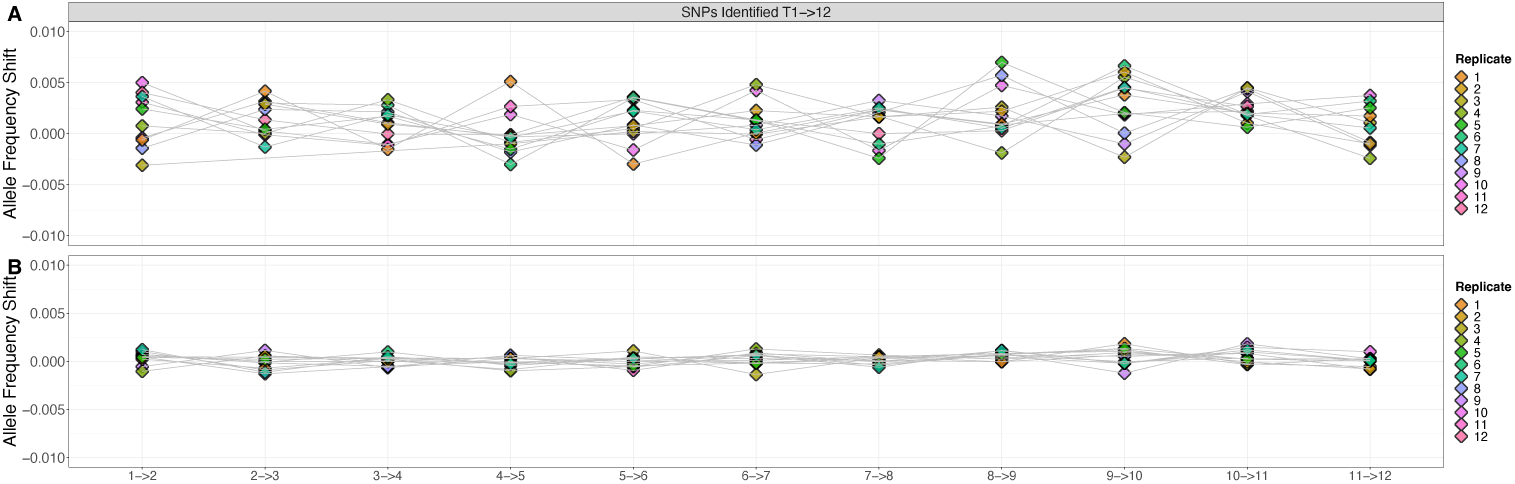
| Median shifts at target SNPs (A) and matched control SNPs (B) identified in each possible eleven time point interval and measured across each pair of single time points. Point color corresponds to replicate cage ID’s and grey lines connect the median shifts for each cage between consecutive time point intervals.

**Figure S23.**
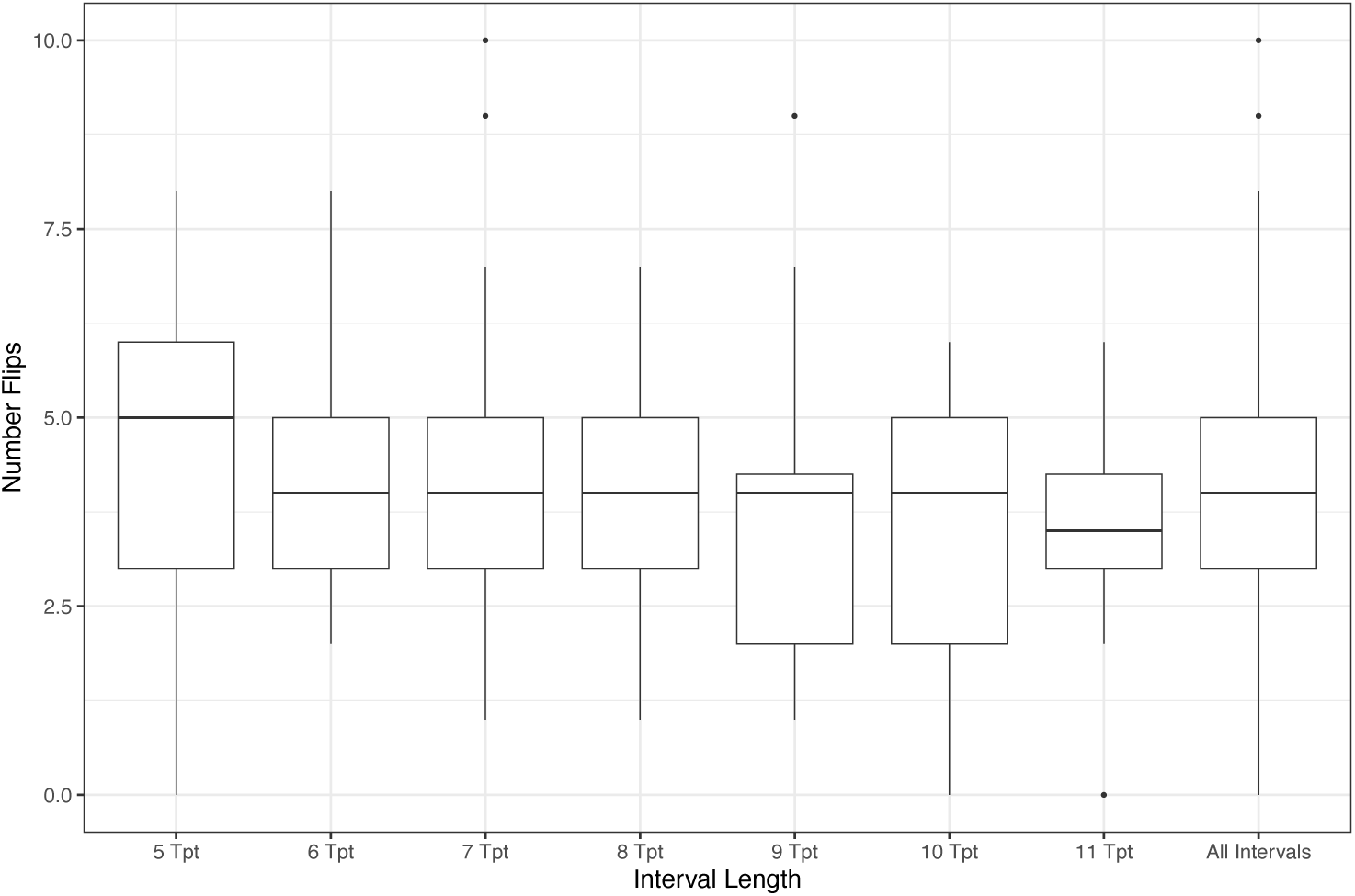
Distributions of the number of flips in the dominant sign of selection between consecutive time point intervals on sets of target SNPs identified across all interval lengths and comparisons and tested in left-out replicate cages. A flip in the dominant sign of selection was inferred if, for an individual cage, the distribution of shifts at SNPs changed from significantly greater to significantly less (or vice versa) than that of the matched control shifts across consecutive time point intervals. All sets of test SNPs were generated using a GLM and leave-one-out approach.

**Figure S24.**
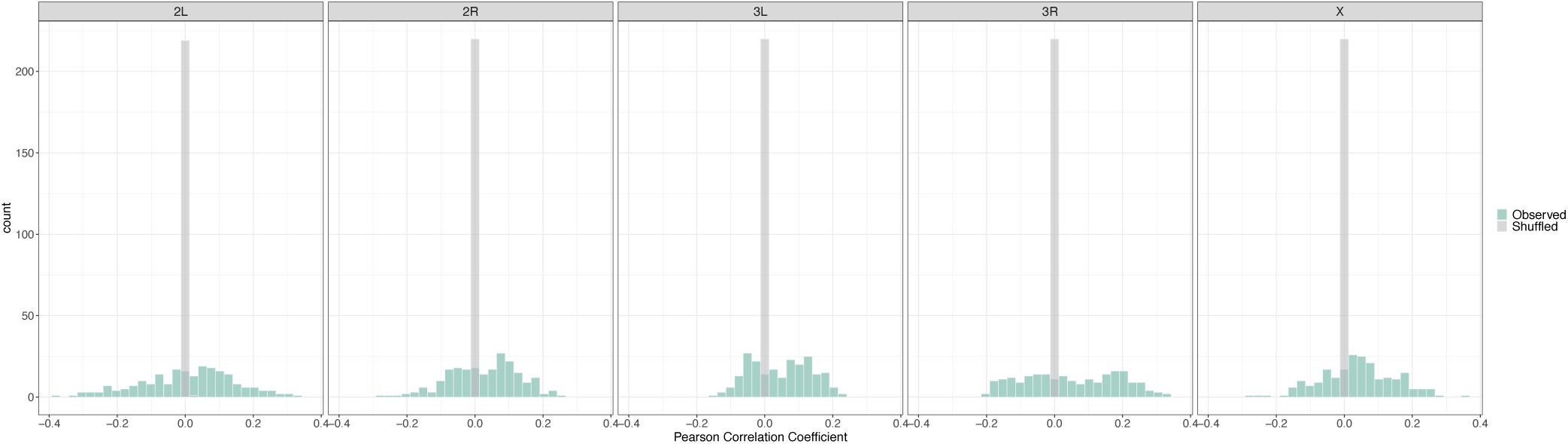
Histogram distributions of Pearson correlations computed between allele frequency shifts for each monthly interval of the 2014 experiment (N = 4), and each interval of the 2021 experiment (N = 56), segregated by chromosomal arm. Bar color correspond to coefficients generated via site information permutations (grey) and observed values (green).

